# Single molecule tracking of bacterial cell surface cytochromes reveals dynamics that impact long-distance electron transport

**DOI:** 10.1101/2021.11.02.466829

**Authors:** Grace W. Chong, Sahand Pirbadian, Yunke Zhao, Lori A. Zacharoff, Fabien Pinaud, Mohamed Y. El-Naggar

## Abstract

Using a series of multiheme cytochromes, the metal-reducing bacterium *Shewanella oneidensis* MR-1 can perform extracellular electron transfer (EET) to respire redox-active surfaces, including minerals and electrodes outside the cell. While the role of multiheme cytochromes in transporting electrons across the cell wall is well established, these cytochromes were also recently found to facilitate long-distance (micrometer-scale) redox conduction along outer membranes and across multiple cells bridging electrodes. Recent studies proposed that long-distance conduction arises from the interplay of electron hopping and cytochrome diffusion, which allows collisions and electron exchange between cytochromes along membranes. However, the diffusive dynamics of the multiheme cytochromes have never been observed or quantified *in vivo*, making it difficult to assess their hypothesized contribution to the collision-exchange mechanism. Here we use quantum dot labeling, total internal reflection fluorescence microscopy, and single-particle tracking to quantify the lateral diffusive dynamics of the outer membrane-associated decaheme cytochromes MtrC and OmcA, two key components of EET in *S. oneidensis*. We observe confined diffusion behavior for both quantum dot-labeled MtrC and OmcA along cell surfaces (diffusion coefficients *D*_MtrC_ = 0.0192 ± 0.0018 μm^2^/s, *D*_OmcA_ = 0.0125 ± 0.0024 μm^2^/s) and the membrane extensions thought to function as bacterial nanowires. We find that these dynamics can trace a path for electron transport via overlap of cytochrome trajectories, consistent with the long-distance conduction mechanism. The measured dynamics inform kinetic Monte Carlo simulations that combine direct electron hopping and redox molecule diffusion, revealing significant electron transport rates along cells and membrane nanowires.

**Significance:** Multiheme cytochromes in *Shewanella oneidensis* MR-1 transport electrons across the cell wall, in a process called extracellular electron transfer. These electron conduits can also enable electron transport along and between cells. While the underlying mechanism is thought to involve a combination of electron hopping and lateral diffusion of cytochromes along membranes, these diffusive dynamics have never been observed *in vivo*. Here, we observe the mobility of quantum dot-labeled cytochromes on living cell surfaces and membrane nanowires, quantify their diffusion with single-particle tracking techniques, and simulate the contribution of these dynamics to electron transport. This work reveals the impact of redox molecule dynamics on bacterial electron transport, with implications for understanding and harnessing this process in the environment and bioelectronics.

## Introduction

Redox reactions are a fundamental part of how many organisms extract energy for life; this process involves the transfer of electrons from an electron donor to an electron acceptor, through the cellular electron transport chain (1). *Shewanella oneidensis* MR-1 is a Gram-negative, facultative anaerobic bacterium that can gain energy by utilizing a diverse array of electron acceptors, from soluble molecules like oxygen to insoluble objects outside the cell surface, including minerals and electrodes (2). This respiratory versatility is made possible by a series of multiheme *c*-type cytochromes that transport electrons from the electron transport chain on the inner membrane, across the otherwise electrically insulating periplasmic space and outer membrane, to solid materials outside the cell, in a process known as extracellular electron transfer (EET) (2–4). The capability to perform EET makes *S. oneidensis* and other electroactive microorganisms particularly interesting for applications in bioelectrochemical technologies, such as microbial fuel cells and microbial electrosynthesis (3, 5, 6), as well as emerging concepts for living electronics (7). Since the discovery of *S. oneidensis* (8), extensive studies have revealed an EET network of cytochromes that bridge the cell envelope, including inner membrane tetraheme cytochrome CymA, periplasmic cytochromes such as STC, outer membrane porin-cytochrome complexes such as MtrAB and cell surface cytochromes such as MtrC and OmcA (4, 9, 10). MtrA is a decaheme cytochrome located on periplasmic side of the outer membrane and connected to the surface by the transmembrane porin MtrB, where the outwardmost heme in MtrA can then interact with cell surface cytochromes such as MtrC and OmcA (9, 10), which can in turn act as an external interface between the cell and extracellular electron acceptors (4). The decaheme outer membrane-associated MtrC and OmcA in particular are largely extracellularly exposed (10), attached to the cell surface by a lipidated cysteine at the N-terminus (11). Once electrons have reached the cell surface, they can be transferred to extracellular electron acceptors by direct contact with these cell surface cytochromes or indirectly via soluble redox shuttles, such as flavins (3).

In addition to bridging the gap between the electron transport chain on the inner membrane and electron acceptors outside the cell, EET components can also enable long-distance (micrometer scale) lateral electron transport along membranes and across multiple cells, as demonstrated recently with electrochemical gating measurements of electron conduction through *S. oneidensis* cellular monolayers connecting interdigitated electrodes (12). This multi-cell redox conduction process is dependent on the presence of the Mtr/Omc EET pathway cytochromes, and exhibits a thermal activation energy consistent with the activation barrier for transport through the decaheme chain of MtrC (12, 13). These observations, and other remarkable demonstrations of cytochrome-mediated redox conduction in electroactive biofilms (14), motivate a better understanding of the cytochrome density and physical electron transport mechanism that can give rise to long-distance conduction. Previous measurements estimate high densities of MtrC and OmcA on the *S. oneidensis* cell surface (up to 30,000 proteins/μm^2^) (15), but this surface coverage is not sufficient to provide a crystalline-like packing that allows direct inter-protein electron hopping along the full micrometer-scale conduction path. Additional knowledge about the cytochrome distribution was recently provided by electron cryotomography (ECT) of the *S. oneidensis* outer membrane extensions (16). These extensions, proposed to function as nanowires for electron transport, contain the EET components, are known to form after cell-surface attachment, and have been observed up to 100 μm in length at an elongation rate of 40 μm/h (17, 18). They have been observed with a range of morphologies, from tubule-like structures to vesicle chains (16, 17, 19). The ECT observations revealed a heterogeneous distribution of outer membrane-associated cytochromes, with inter-protein spacings ranging from immediately adjacent to being separated by tens of nanometers (16). In light of these findings, we previously proposed a collision-exchange model (Fig. 1A), where the lateral diffusion of the multiheme cytochromes leads to collisions and inter-protein electron exchange along the membrane (16). This mechanism, which accounts for both direct electron hopping between redox centers and their physical diffusion, played a critical role in understanding conduction through redox polymers (20), but remains underexplored in the context of EET. Recent studies have indeed hinted at the importance of cytochrome mobility as a possible contributor to long-distance conduction (12, 16, 21, 22), but this contribution has not been verified with experimental measurements of the diffusive dynamics of the membrane-associated EET cytochromes *in vivo*.

**Fig. 1.**
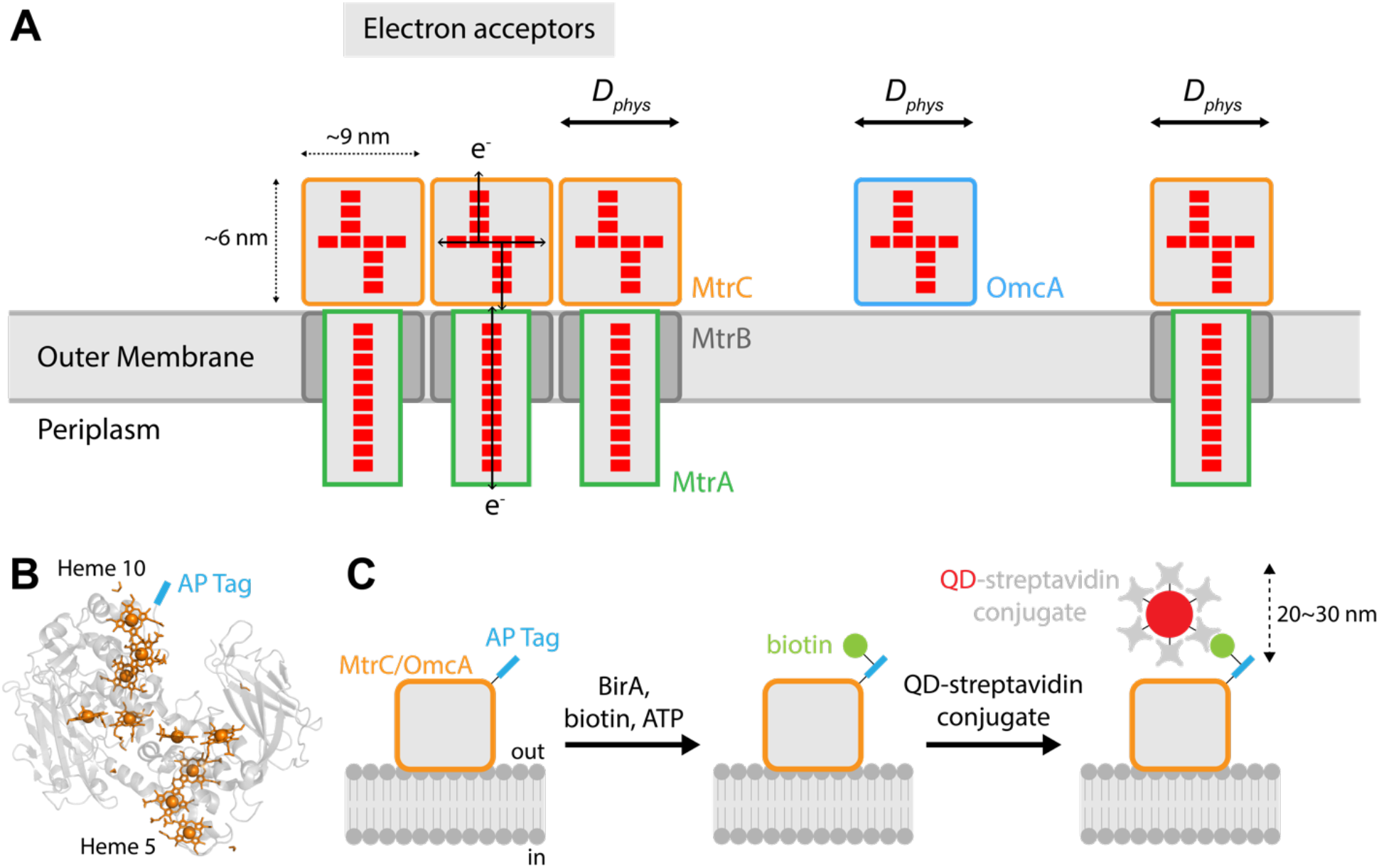
Lateral diffusion and labeling strategy. (*A*) Schematic of diffusion-assisted electron hopping along the *Shewanella oneidensis* MR-1 outer membrane. Lateral motion of multiheme cytochromes (diffusion coefficient *D*_*phys*_) leads a collision-exchange mechanism of inter-protein electron transport over large distances. Red spots represent hemes in multiheme cytochromes. Labeled proteins are outer membrane cytochromes MtrC and OmcA, outer membrane-associated periplasmic cytochrome MtrA, and outer membrane porin MtrB. (*B*) Structure of MtrC (PDB ID 4LM8) illustrates location of biotin acceptor peptide (AP) tag, fused to C-terminus of MtrC (or OmcA) near Heme 10. Hemes and porphyrin rings are colored orange, and AP tag is colored blue. (*C*) Schematic of labeling strategy, adapted from (27). The biotin acceptor peptide (AP: GLNDIFEAQKIEWHE) is fused to MtrC (or OmcA). At the cell surface, biotin ligase BirA biotinylates the AP, and QD-streptavidin conjugates bind the biotinylated MtrC-AP (or OmcA-AP).

While it is expected that membrane components are capable of diffusion, it turns out that such dynamics have rarely been measured for outer membrane proteins in Gram-negative bacteria. In fact, few unique outer membrane proteins have actually been studied, and most existing studies generally involve β-barrel porin or channel-type integral membrane proteins in *Escherichia coli* (23–25). Out of several commonly used methods for studying diffusion dynamics, single-particle tracking (SPT) combined with total internal reflection fluorescence (TIRF) microscopy provides several advantages, such as high spatial resolution that enables precise localization and tracking of individually labeled proteins (26). The absence of experimental data on the mobility of EET components motivated us to apply these single-molecule techniques to assess their dynamics on the outer membrane of *S. oneidensis*. To our knowledge, this work presents the first measurements of the diffusive dynamics of bacterial extracellular electron conduits, and it adds to the relatively short list of cell surface proteins whose diffusion was measured in prokaryotes.

In this study, we set out to (*i*) visualize individual cell surface cytochromes on living cells, (*ii*) assess their mobility along the membrane surfaces, (*iii*) quantify their diffusive dynamics, and (*iv*) investigate how this diffusion impacts overall electron transport in the context of the collision-exchange mechanism. To observe the dynamics of individual cytochromes, we tagged the *S. oneidensis* cell surface cytochromes MtrC and OmcA with biotin acceptor peptides, which allowed site-specific targeting of the biotinylated proteins with streptavidin-conjugated quantum dots. This labelling strategy allowed us to measure the diffusive dynamics of the quantum dot labelled electron conduits with *in vivo* TIRF microscopy and single-particle tracking (27–30). We then quantified the diffusion coefficients of both MtrC and OmcA along the surface of the outer membrane and membrane nanowires and calculated the contribution of these dynamics to long-distance electron conduction through kinetic Monte Carlo simulations. Altogether, our study suggests that the dynamics of EET components play an important role in overall electron transport over micrometer length scales.

## Results and Discussion

### Successful and specific *in vivo* labeling of cell surface cytochromes MtrC and OmcA

We used the labeling scheme described in (27, 28, 31) to label cell surface cytochromes MtrC and OmcA in *S. oneidenis* MR-1. Briefly, as pictured in Fig. 1B-C, a 15-amino acid biotin acceptor peptide (AP) tag from *E. coli* (32) was fused to the C-termini of MtrC and OmcA. Once assembled in the periplasm and exported to the outer membrane, cytochromes expressing the AP tag can then be biotinylated externally by the addition of biotin ligase BirA. Finally, the biotinylated cytochromes can then be detected by streptavidin-conjugated probes, which would allow the labeled cytochromes to be imaged in real-time by microscopy. Another benefit of this labeling scheme, which combines a small peptide tag with extracellular labeling (Fig. 1C), is to minimize interference to the localization of MtrC and OmcA on the outer surface of the cell, where peptides produced in the cytoplasm are transported to the periplasm for protein folding and heme assembly before being exported to the extracellular side of the outer membrane (33).

DNA inserts for AP-tagged MtrC and OmcA (SI Appendix, Fig. S1 and S2) were constructed by overhang PCR and cloned into the pBBR1-MCS2 plasmid (34). Plasmid constructs were transformed into respective *S. oneidensis* MR-1 Δ*mtrC* or Δ*omcA* deletion backgrounds from (35). All strains, plasmids, and primers used in this study are listed in Table 1 and SI Appendix, Table S1. Cytochrome presence was then detected by staining SDS-PAGE gels with a heme-reactive peroxidase activity assay using 3,3’-diaminobenzidine and hydrogen peroxide (SI Appendix, Fig. S3), which confirmed that the AP-tagged strains produced heme-containing proteins of expected size, compared to positive and negative controls (wild type and gene deletion mutant). Sanger sequencing of plasmids purified from final host strains also verified sequence integrity of the AP tag.

**Table 1.**
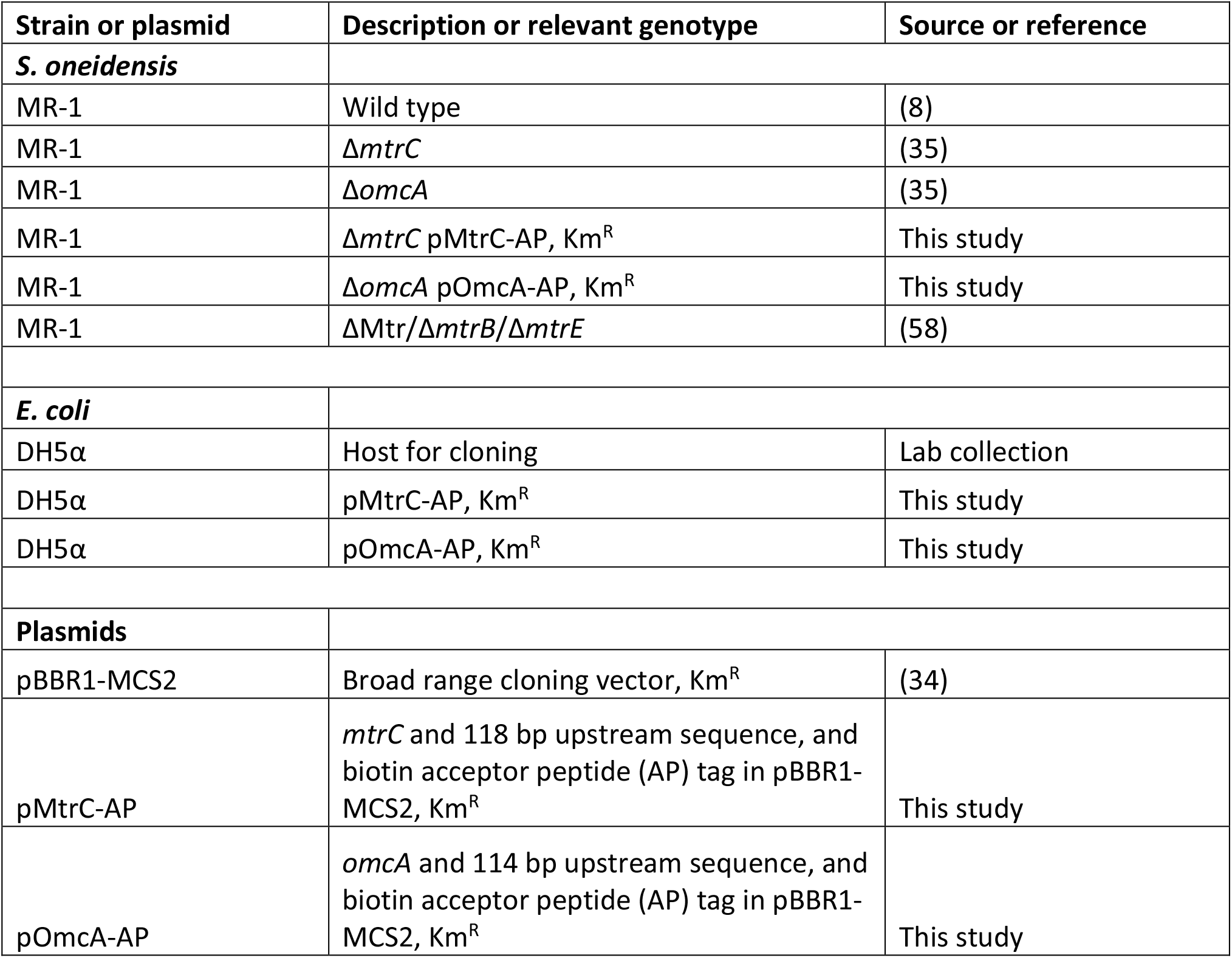
Strains and plasmids used in this study.

Next, we performed Western blot and microscopy controls where we systematically omitted key components in the labeling process, to confirm that the labeling scheme works in our system (Fig. 2 and SI Appendix, Fig. S4). In Western blots probing for biotinylated proteins using streptavidin-horseradish peroxidase, MtrC-AP (or OmcA-AP) were detected only when all key components of the labeling process were provided (Fig. 2A and SI Appendix, Fig. S4A), indicating that the tagged protein was successfully and specifically biotinylated and detected by the streptavidin probe. Similarly, microscopy labeling controls were performed, where biotinylated proteins were visualized by streptavidin-conjugated Alexa Fluor 647 (Fig. 2B and SI Appendix, Fig. S4B). Though cells were visible by standard brightfield imaging in all samples, strong fluorescent signal visualizing biotinylated proteins were only detected in the condition where all key labeling components were present. Taken collectively, these Western blot, fluorescence microscopy, and associated controls demonstrate successful and specific labeling of MtrC and OmcA. Furthermore, our ability to perform extracellular *in vivo* labeling and subsequent microscopic detection of MtrC and OmcA via a C-terminal AP tag is consistent with the recently published orientation of MtrC relative to the MtrAB transmembrane complex (10), where Heme 10 (C-terminal side) is extracellularly exposed and Heme 5 (N-terminal side) is facing the cell surface.

**Fig. 2.**
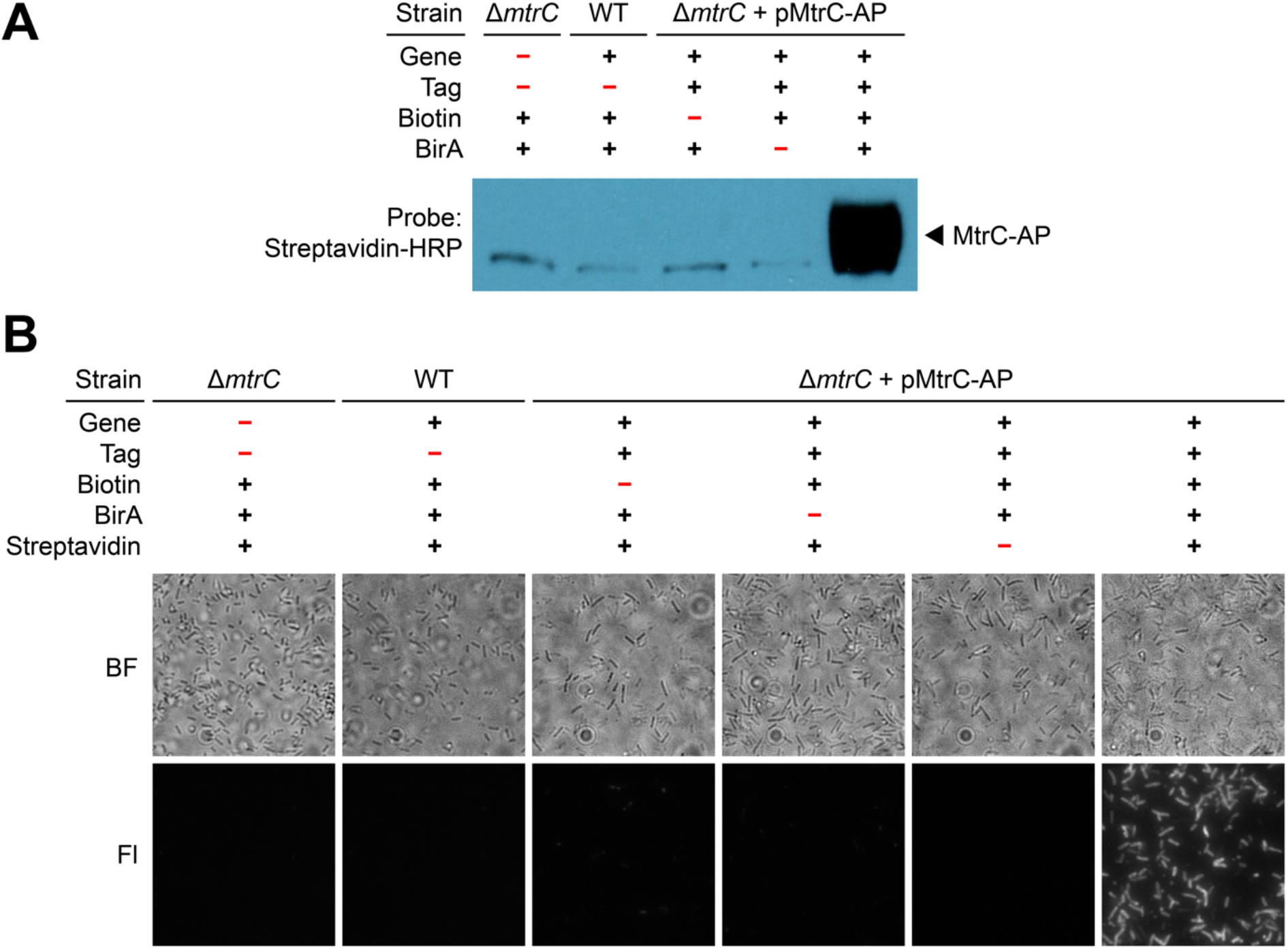
Key labeling controls demonstrate successful and specific labeling of MtrC. (*A*) Western blot labeling control for MtrC where key parts of the labeling process were systematically omitted. When using streptavidin (streptavidin-horseradish peroxidase, HRP) to probe for biotinylated proteins, a thick dark band of biotinylated MtrC-AP is detected only in Lane 5 when all key components are present. The faint band slightly below labeled MtrC-AP (approx. 79.6 kDa) and present in all samples is an endogenously biotinylated protein (acetyl-CoA carboxylase, approx. 76 kDa) (57). (*B*) Microscopy labeling control for MtrC where key parts of the labeling process were systematically omitted. Top row contains brightfield (BF) images showing many cells in each sample. Bottom row images show fluorescence (Fl) signal from streptavidin-conjugated Alexa Fluor 647 that was used to detect biotinylated MtrC-AP; fluorescence labeling was detected strongly in the bottom right image, and only when all key labeling components were present. All microscopy images are 36.5 μm by 36.5 μm.

### Single-particle imaging and tracking reveals mobility of MtrC and OmcA along cell surface and membrane extensions

Once the labeling scheme was established in our system, we investigated cell surface protein dynamics using targeted quantum dot (QD) labeling and single-particle tracking (SPT) (24, 30, 36). QDs were chosen as the fluorescent label due to their high signal-to-noise ratio and photostability, which makes them useful for SPT (30, 36). In addition, this labeling scheme takes advantage of the very strong (femtomolar scale) binding affinity and very low dissociation rate between biotin and streptavidin, which makes biotin-streptavidin labeling schemes useful for single molecule labeling and other applications (37, 38). To test the hypothesis that MtrC and OmcA are mobile along the cell surface and to quantify their diffusion behavior, we labeled MtrC-AP and OmcA-AP by *in vivo* biotinylation and streptavidin-conjugated QDs (Fig. 1B-C) and imaged their dynamics on the surface of living cells by dual-color time-lapse total internal reflection fluorescence (TIRF) microscopy. To visualize the cell outer membrane and membrane extensions, we used FM 1-43FX, a lipid membrane dye. To visualize individual cytochromes, we titrated the concentration of streptavidin-conjugated QDs until it was possible to distinguish individual particles (e.g. 1-5 QDs/cell).

We observed that MtrC and OmcA are indeed mobile along the cell surface and outer membrane extensions, and we traced their mobility with SPT (Fig. 3 and Movies S1-S3). Briefly, SPT detects the position of each QD molecule in each frame, and connects these detected positions frame-by-frame to build trajectories over time. In our experiments, the typical QD localization precision was ∼15 nm. Fig. 3 highlights the workflow of single QD detection and tracking as applied to labeling of OmcA on the *S. oneidensis* cell surface. Starting with a large field of view (Fig. 3A), the fluorescence of individual QDs across hundreds of cells was tracked, typically over 1-2 min, with an acquisition rate of 40 ms/frame to generate thousands of trajectories of cytochrome diffusion. Individual QD trajectories were typically punctuated by gaps resulting from the expected blinking behavior of single QD molecules (39). Zooming in on individual cells (Fig. 3B-C, corresponding to dashed areas in Fig. 3A) highlights the heterogeneous behavior of diffusing cytochromes, which was further analyzed to classify and quantify diffusive dynamics.

**Fig. 3.**
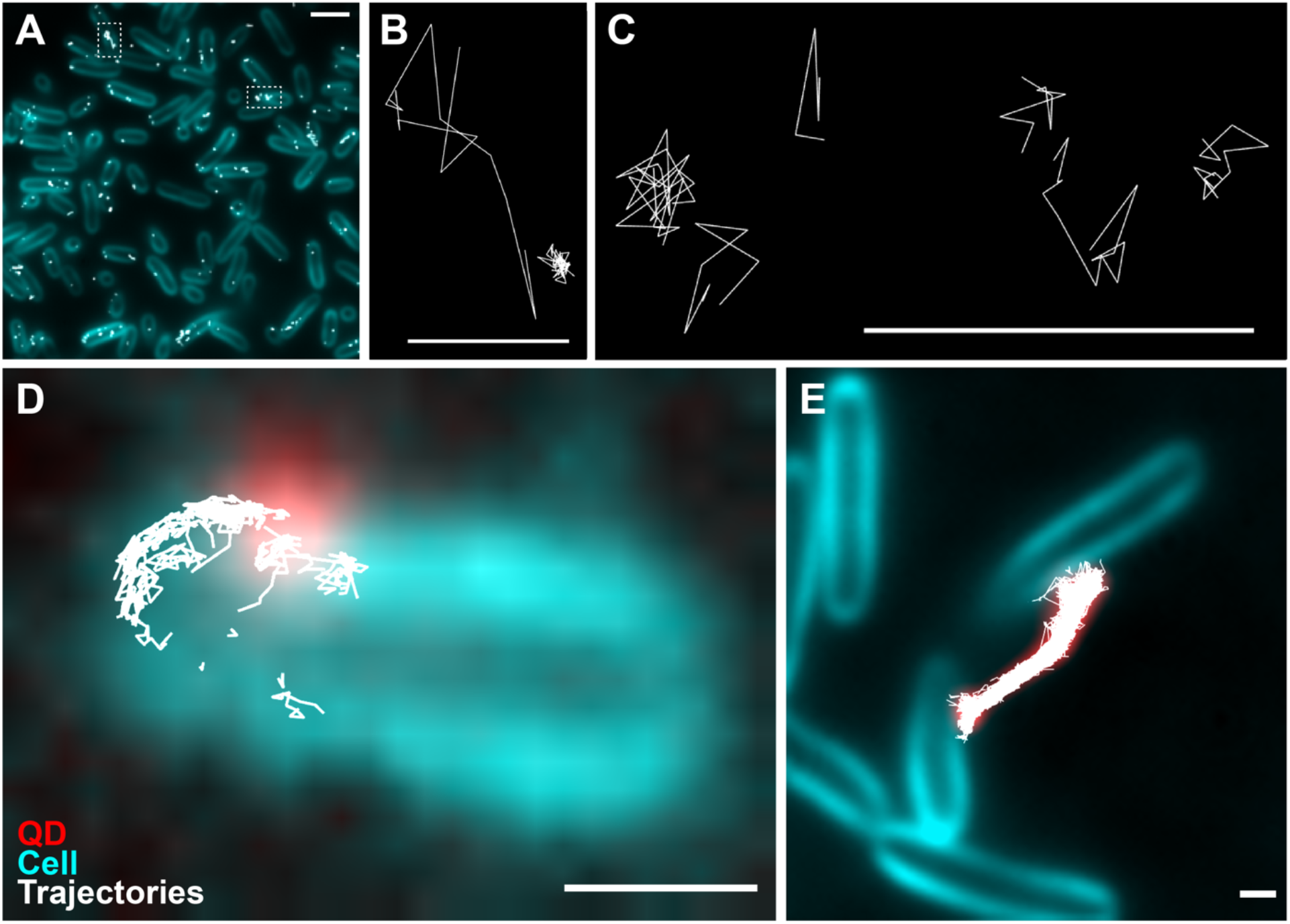
Imaging and single molecule tracking of quantum dot (QD)-labeled OmcA using total internal reflection fluorescence (TIRF) microscopy. (*A*) Snapshot of OmcA-AP trajectories (white) in multiple cells (cyan). Trajectories from 1.5 min of time-lapse microscopy (40 ms/frame) were overlaid onto the corresponding mean intensity projection image of cells labeled with lipid membrane dye FM 1-43FX. Trajectories in white dashed boxes are blown up in panels B and C. Scale bar: 2 μm. (*B-C*) Some example trajectories from the two cells outlined in panel A, ranging from 0.16-1.16 s in duration. Scale bars: 500 nm. *(D-E)* Streptavidin-conjugated QD705 was used to detect exogenously biotinylated OmcA-AP (red). Cell membrane and membrane extensions are labeled by FM 1-43FX (cyan). (*D*) Trajectories from a single QD-labeled OmcA-AP as it moved along the surface of a cell, as seen in Movie S1. Here, the QD signal (red) and its trajectories (white) are overlaid with the mean intensity projection image of the cell (cyan). For clarity, only the first frame of QD signal is shown; trajectories are from the entire video (86 s, 40 ms/frame). Scale bar: 500 nm. (*E*) Snapshot of OmcA-AP trajectories overlaid on an outer membrane extension. Trajectories (white) are from 6 min (40 ms/frame) of time-lapse microscopy tracing several QD-labeled OmcA-AP (red; mean intensity projection image) on a membrane extension that appears to connect two cells (cyan; mean intensity projection image). A short portion (12 s) of the time-lapse corresponding to this panel can be seen in Movie S2. Scale bar: 500 nm.

When viewed in the context of the cell surface (Fig. 3D) and membrane extensions (Fig. 3E), we observed that QD-labeled MtrC and OmcA can both explore a significant fraction of the underlying membrane surface. In addition, we observed significant overlap in diffusion trajectories for multiple cytochromes, notably along a membrane extension linking two cells shown in Fig. 3E. These observations support our proposed collision-exchange mechanism for long-distance electron conduction (Fig. 1A) (16), where diffusive dynamics can bridge gaps between cytochromes and, combined with direct electron hopping, lead to a continuous path for electron transport along the membrane. While long-distance multi-cell conduction was recently observed by electrochemical gating, and cytochrome diffusion was proposed to play a role (12), our measurements provide the first direct look at these dynamics. Next, we sought to quantitatively analyze the diffusion characteristics in order to assess their contribution to biological electron transport over micrometer length scales.

### Quantifying the dynamics of MtrC and OmcA along the cell surface and membrane extensions

To quantify the observed cytochrome mobility, we performed diffusion analyses for MtrC or OmcA trajectories diffusing either on the cell surface or on membrane extensions. Methods for SPT and determination of diffusion coefficients have previously been described in detail (26, 30, 40–42). All diffusion coefficients determined in this study are listed in Table 2 and described below.

**Table 2.** Summary of diffusion coefficients (*D*) and confinement radii (*R*) determined in this study using a 1-component or 2-component model of diffusion (Fig. 4-7). Percentages indicate the fraction of all trajectories belonging to each respective component. *D* and *R* values were determined according to a confined diffusion model (Eqn. 2).

| Protein | Surface | Fraction | $D$ ($\mu\text{m}^2/\text{s}$) | $R$ (nm) |
| --- | --- | --- | --- | --- |
| **MtrC** | **Cell** | **100%** | **$0.0192 \pm 0.0018$** | **80** |
| *MtrC1* (Slow component) | Cell | 90% | $0.00235 \pm 0.00112$ | 19 |
| *MtrC2* (Fast component) | Cell | 10% | $0.124 \pm 0.010$ | 264 |
| **OmcA** | **Cell** | **100%** | **$0.0125 \pm 0.0024$** | **59** |
| *OmcA1* (Slow component) | Cell | 94% | $0.000577 \pm 0.000152$ | 18 |
| *OmcA2* (Fast component) | Cell | 6% | $0.0939 \pm 0.0059$ | 242 |
| **MtrC** | **OME** | **100%** | **$0.00945 \pm 0.00028$** | **132** |
| *MtrC1* (Slow component) | OME | 66% | $0.00162 \pm 0.00011$ | 52 |
| *MtrC2* (Fast component) | OME | 34% | $0.0353 \pm 0.0021$ | 198 |
| **OmcA** | **OME** | **100%** | **$0.0102 \pm 0.0002$** | **112** |
| *OmcA1* (Slow component) | OME | 82% | $0.00188 \pm 0.00006$ | 46 |
| *OmcA2* (Fast component) | OME | 18% | $0.0482 \pm 0.0016$ | 242 |

First, we evaluated the general diffusion of MtrC or OmcA on the cell surface by pooling data from all trajectories in each dataset and constructing ensemble mean squared displacement (MSD) curves (Fig. 4). We found that both MtrC and OmcA exhibit confined diffusion behaviors, with MSD curves reaching a plateau over timescales < 1 s. By fitting these curves with a confined diffusion model (SI Appendix, *Materials and Methods*), we determined the overall diffusion coefficients *D* and confinement radii *R* for QD-labeled MtrC (*D* = 0.0192 ± 0.0018 μm^2^/s; *R* = 80 nm) and OmcA (*D* = 0.0125 ± 0.0024 μm^2^/s, *R* = 59 nm). While quantitative information regarding the diffusion of prokaryotic cell surface proteins is limited, our measurements of MtrC and OmcA (Table 2) are consistent in magnitude with the observations made for other bacterial outer membrane proteins, which are on the scale of *D* = 0.006-0.15 μm^2^/s and *R* = 15-300 nm (23–25, 43). Moreover, the confinement radii of MtrC and OmcA are consistent with our previous measurements of center-to-center distances between putative cell surface cytochromes in *S. oneidensis* (16). We note that while a single cytochrome does not typically travel out of a confinement domain, multiple cytochromes might populate, diffuse, and collide within the same region. Proteins can also stochastically escape an area of confinement and diffuse more freely across a larger distance of the cell surface over time, before encountering other obstacles.

**Fig. 4.**
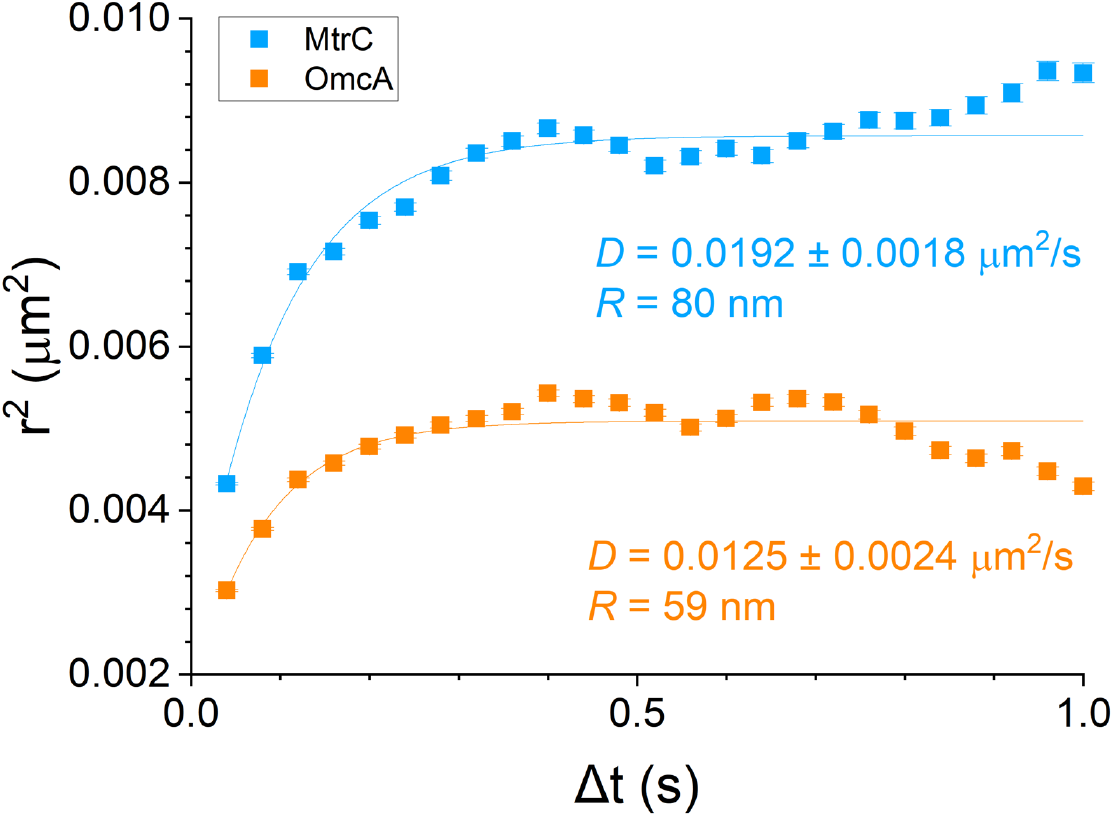
Ensemble mean squared displacement (MSD) analysis shows overall confined diffusion behavior by MtrC (blue) and OmcA (orange) on the cell surface. Y-axis shows mean displacement squared (*r*^*2*^) for each time lag (*Δt*) on the X-axis. Fitting the plots with a confined diffusion model (Eqn. 2) yields diffusion coefficients *D* and confinement radii *R* as labeled. Error bars show 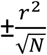 where *N* is the number of independent data points (i.e., displacements) analyzed for a given *Δt*. These curves represent 7,678 MtrC-AP and 7,109 OmcA-AP trajectories on the cell surface, from 500-1,000 cells each.

To estimate the smallest diffusion coefficient measurable under our experimental conditions, we also quantified the apparent diffusion of QDs imaged on coverslips, under cell-free conditions. Unsurprisingly, we observe much slower mobility, with *D* = 3.56 × 10^−5^ ± 1.62 × 10^−5^ μm^2^/s (SI Appendix, Fig. S5), clearly distinct from the lateral diffusion of MtrC and OmcA (Fig. 4, Table 2). To rule out the possibility that the observed cytochrome diffusion is influenced by streptavidin-conjugated QDs binding multiple targets (30), SPT was also performed in the presence of excess free biotin, added to cells immediately after QD labeling in order to saturate residual streptavidin binding sites. Under biotin saturation, no change in the distribution of diffusion coefficients was observed (SI Appendix, Fig. S6), indicating that the membrane mobility of MtrC and OmcA is not impacted by cross-linking from multivalent streptavidin QDs.

As seen in Fig. 3, heterogeneities in the shape of diffusing trajectories (e.g. Fig. 3D, Movie S1) suggest that MtrC and OmcA might transition between multiple diffusing behaviors. To address this possibility, the ensemble of trajectories for each type of cytochrome was analyzed by probability distribution of square displacements (PDSD) (40, 42, 44) (Fig. 5). We found that the diffusion of MtrC could be described by a 2-component model, with 90% of MtrC displaying a slow and confined mobility with diffusion coefficient *D*_*1*_ = 0.00235 ± 0.00112 μm^2^/s and confinement radius *R*_*1*_ = 19 nm, while a 10% minority diffuses significantly faster over less confined membrane regions (*D*_*2*_ = 0.124 ± 0.010 μm^2^/s, *R*_*2*_ = 264 nm). Likewise, the majority of OmcA displayed a slow and confined diffusive behavior (94%, *D*_*1*_ = 0.000577 ± 0.000152 μm^2^/s, *R*_*1*_ = 18 nm) together with a less prevalent but faster diffusion over large membrane domains (6%, *D*_*2*_ = 0.0939 ± 0.0059 μm^2^/s, *R*_*2*_ = 242 nm). The faster, less confined diffusion detected by this analysis (Fig. 5B,D) may represent events where generally confined redox proteins escape crowded areas and diffuse more freely across the bacterial membrane, their diffusion being limited by the overall size of the cell itself, typically 500 nm in diameter. The detection of two diffusive behaviors for MtrC and OmcA is also consistent with the heterogeneity in distribution of proteins along the cell surface previously observed by electron cryotomography (16), which is a common feature among membrane proteins in bacteria (25, 45).

**Fig. 5.**
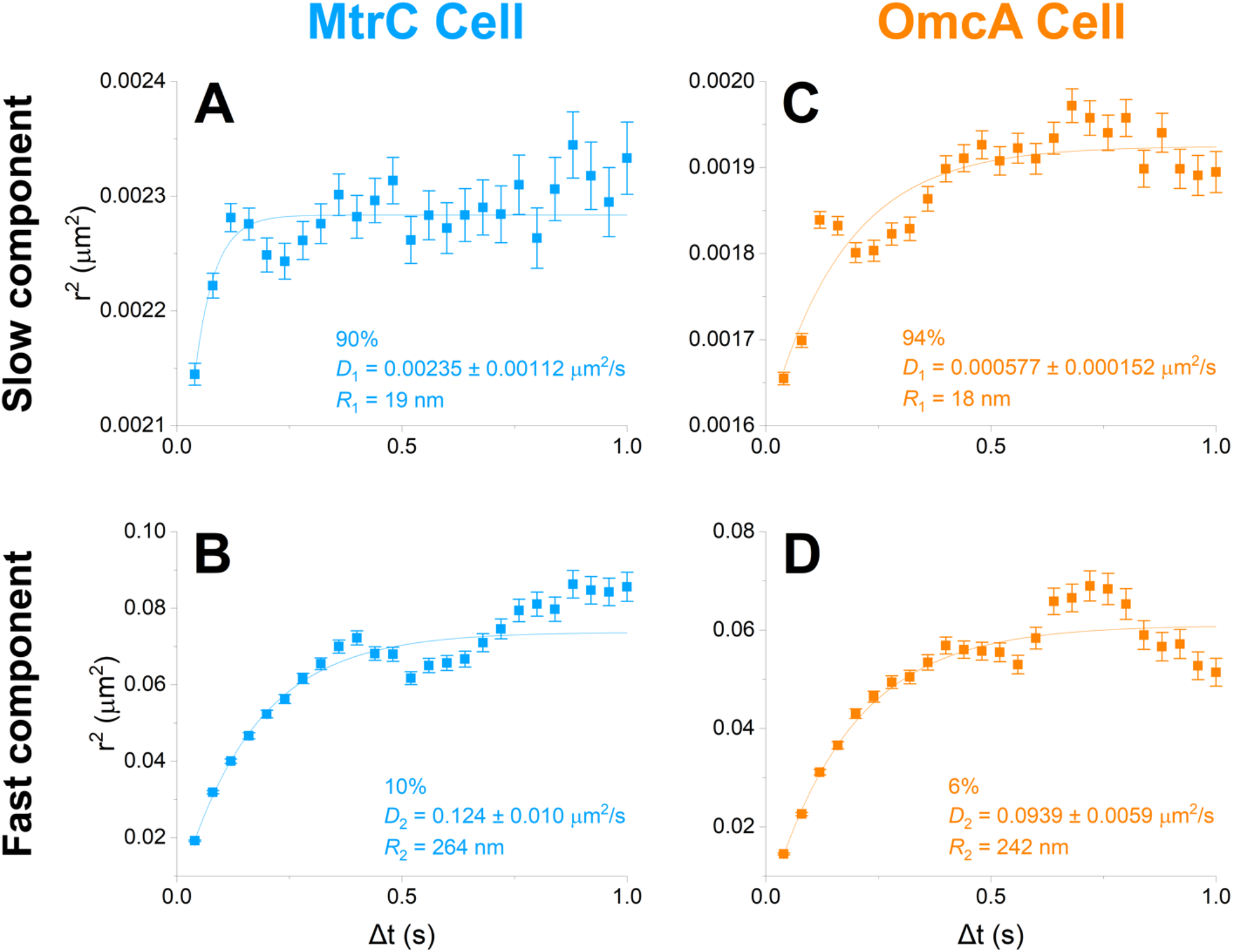
Diffusion analyses for MtrC (blue) and OmcA (orange) on the cell surface, using a 2-component model of diffusing behavior. Left (*A,B*): MtrC-AP. Right (*C,D*): OmcA-AP. On the cell surface, MtrC and OmcA diffusion can be described by two behaviors: (*A,C*) a slower, more confined majority and (*B,D*) a faster, less confined minority. Percentages indicate the respective fractions belonging to each component as determined by probability distribution of square displacement (PDSD) analysis, as described in (40, 42). Ensemble mean squared displacement (MSD) curves were plotted as mean displacement squared *r*^*2*^ as a function of time lag *Δt*. Fitting these curves with a confined diffusion model (Eqn. 2) yields diffusion coefficients *D* and confinement radii *R*, as labeled on each plot. Error bars show 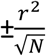 where *N* is the number of independent data points (i.e., displacements) analyzed for each component for a given *Δt*. These curves represent 7,678 MtrC-AP and 7,109 OmcA-AP trajectories on the cell surface, from 500-1,000 cells each.

Next, we investigated the dynamics of MtrC and OmcA on outer membrane extensions (OMEs) of *S. oneidensis*. Compared to cell surface measurements, imaging of QD-labeled cytochromes on OMEs presented technical challenges. Our previous work using a perfusion flow imaging platform (16) allowed robust epifluorescence observations of OME production over time by restricting OMEs to the focal plane using laminar media flow, but this flow was not desired during SPT experiments, since it may interfere with measurements of cytochrome movement. Thus, under our TIRF imaging conditions, OMEs frequently moved in and out of the evanescent excitation field, limiting our ability to easily image these structures and to track individual QDs along them. We therefore limited our analyses to non-moving OMEs that were clearly connected to a cell and were labeled with a low density of QDs. To compensate for the reduced number of QD-labeled OMEs that were optimal for tracking compared to SPT on whole cells, and to record a sufficient number of diffusion steps for analysis, we tracked QDs on OMEs over periods of 6 min.

Ensemble MSD analysis for MtrC and OmcA on OMEs revealed membrane mobilities similar to those observed on the cell surface. The overall diffusion coefficient and confinement radius for MtrC were *D* = 0.00945 ± 0.00028 μm^2^/s and *R* = 132 nm, and for OmcA were *D* = 0.0102 ± 0.0002 μm^2^/s and *R* = 112 nm (Fig. 6). Compared to diffusion on the cell surface (Fig. 4), larger confinement radii for both cytochromes indicate that they are less confined on OMEs than on the bacterial surface. Yet, their respective diffusion coefficients remain on the same order of magnitude, with diffusion being reduced by about 2-fold on OMEs compared to the cell surface for MtrC (*p* < 0.0001, Fig. 4, 6, and SI Appendix, Fig. S7). The reduced confinement of cytochromes on OMEs might stem from differences in the degree of molecular crowding between these extensions and the cell surface. The moderately slower dynamics on OMEs may be related to their morphology, since OMEs can present as vesicle chains with possible junction densities that might limit membrane fluidity at each junction (16). Furthermore, diffusion coefficients on both cells and OMEs may be underestimated by an additional 25-50%, as motion on a 3D tubular membrane surface is projected onto a 2D image plane during SPT and analysis (46, 47); this may contribute to moderately slower dynamics on OMEs relative to cells, as this underestimate is greater for smaller tube diameters (47).

**Fig. 6.**
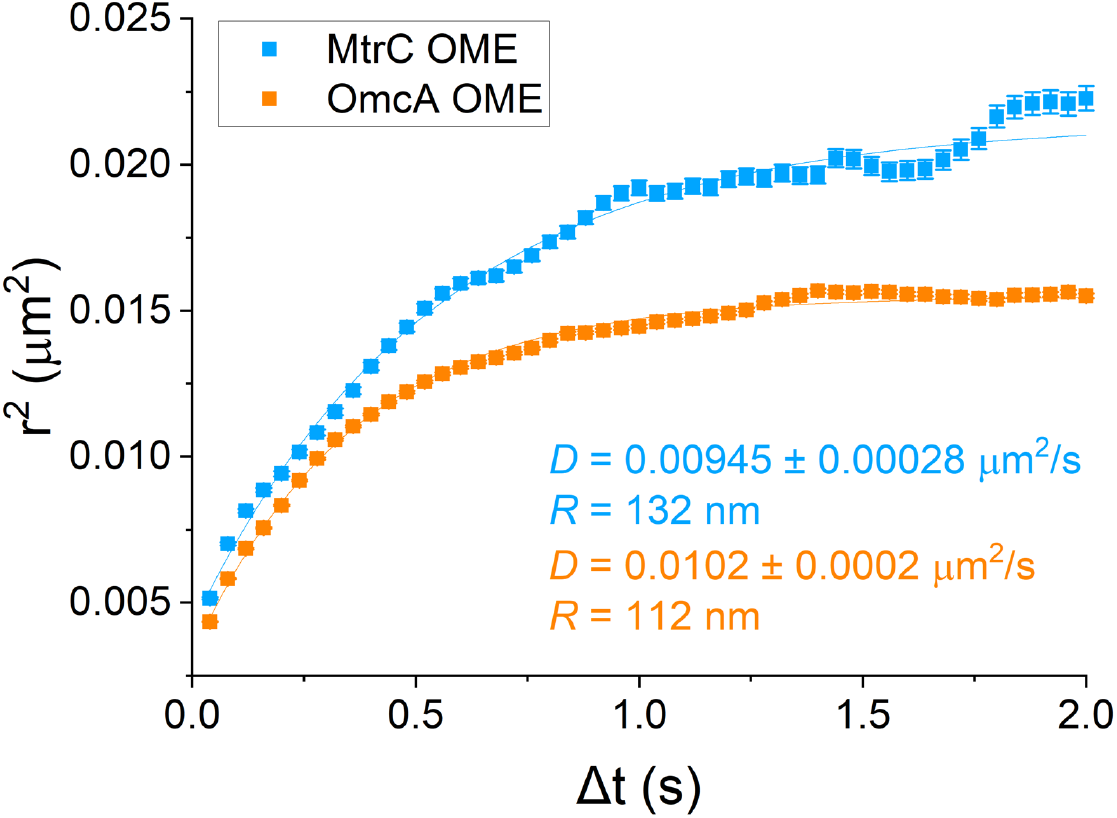
Ensemble mean squared displacement (MSD) analysis shows overall confined diffusion behavior by MtrC (blue) and OmcA (orange) on outer membrane extensions (OMEs). Y-axis shows mean displacement squared (*r*^*2*^) for each time lag (*Δt*) on the X-axis. Fitting the plots with a confined diffusion model (Eqn. 2) yields diffusion coefficients *D* and confinement radii *R* as labeled. Error bars show 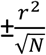 where *N* is the number of independent data points (i.e., displacements) analyzed for a given *Δt*. These curves represent 1,140 MtrC-AP trajectories from 5 OMEs and 5,371 OmcA-AP trajectories from 22 OMEs.

In light of the heterogeneity in diffusion observed on the cell surface (Fig. 5), we also investigated the possibility that MtrC and OmcA exhibit multiple diffusing behaviors on OMEs. Using PDSD analysis, we found that their dynamics on OMEs can also be described by two behaviors: (*i*) a slow and highly confined mobility for a majority of trajectories and (*ii*) a faster and less confined diffusion for a smaller fraction of trajectories (Fig. 7). Diffusion coefficients and confinement radii determined for slow (66%) and fast (34%) MtrC were *D*_*1*_ = 0.00162 ± 0.00011 μm^2^/s, *R*_*1*_ = 52 nm and *D*_*2*_ = 0.0353 ± 0.0021 μm^2^/s, *R*_*2*_ = 198 nm, respectively. Those determined for slow (82%) and fast (18%) OmcA were *D*_*1*_ = 0.00188 ± 0.00006 μm^2^/s, *R*_*1*_ = 46 nm and *D*_*2*_ = 0.0482 ± 0.0016 μm^2^/s, *R*_*2*_ = 242 nm. Generally, both slow and fast MtrC and OmcA had slower mobility on OMEs than their counterparts on the cell surface (Fig. 5, 7, and SI Appendix, Fig. S7B), as predicted by their overall diffusion (Fig. 4, 6, and SI Appendix, Fig. S7A). This is likely due to differences in structure outlined previously. Here, the slow diffusing MtrC and OmcA were noticeably less confined, with over 2.5-fold increase in *R*_*1*_ on OMEs compared to the cell surface. This increase in membrane domain size, from *R*_*1*_ ≈ 19 to 50 nm, may suggest that membrane rearrangement into OMEs allows the highly confined fraction of cytochromes to explore a larger area, their diffusion now being limited by the size of a vesicle/OME itself, approx. 100 nm in diameter.

**Fig. 7.**
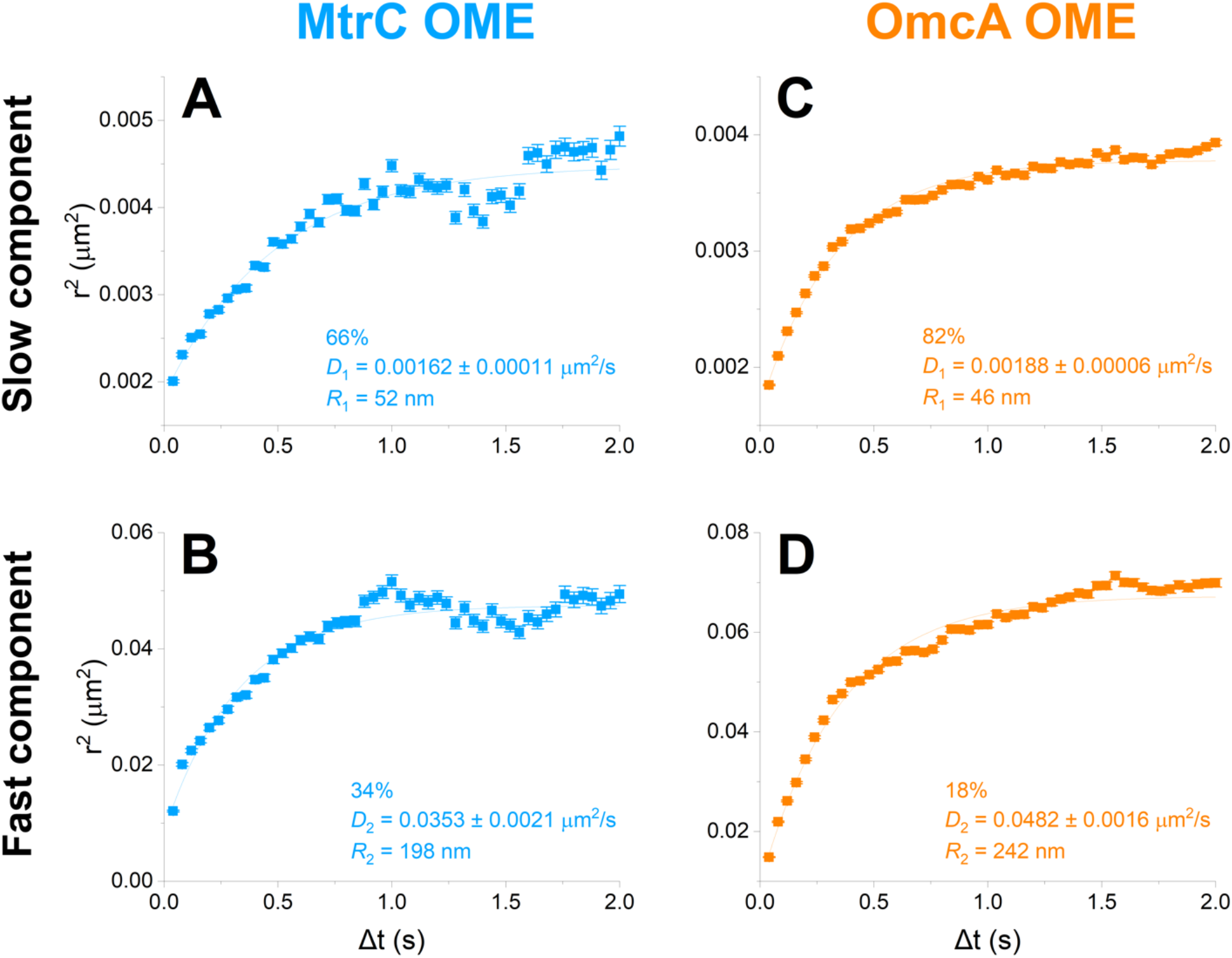
Diffusion analyses for MtrC (blue) and OmcA (orange) on outer membrane extensions (OMEs), using a 2-component model of diffusing behavior. Left (*A,B*): MtrC-AP. Right (*C,D*): OmcA-AP. On OMEs, MtrC and OmcA diffusion can be described by two behaviors: (*A,C*) a slower, more confined majority and (*B,D*) a faster, less confined minority. Percentages indicate the respective fractions belonging to each component as determined by probability distribution of square displacement (PDSD) analysis, as described in (40, 42). Ensemble mean squared displacement (MSD) curves were plotted as mean displacement squared *r*^*2*^ as a function of time lag *Δt*. Fitting these curves with a confined diffusion model (Eqn. 2) yields diffusion coefficients *D* and confinement radii *R*, as labeled on each plot. Error bars show 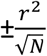 where *N* is the number of independent data points (i.e., displacements) analyzed for each component for a given *Δt*. These curves represent 1,140 MtrC-AP trajectories from 5 OMEs and 5,371 OmcA-AP trajectories from 22 OMEs.

Altogether, MtrC and OmcA appear to display relatively similar diffusive behavior, whether on the cell surface (Fig. 4-5) or on membrane extensions (Fig. 6-7), which is not surprising, since they are structurally and functionally homologous (48). The slightly faster diffusion of MtrC on the cell surface compared to OmcA (+50%, *p* = 0.03, Fig. 4 and SI Appendix, Fig. S7A) could potentially be due to a difference in protein interactions (49, 50).

### Simulations combine electron hopping and cytochrome dynamics to reveal long-distance electron transport along membrane surfaces

To understand the magnitude of long-distance electron conduction that can arise from the interplay of electron hopping and cytochrome motion along membranes, we performed kinetic Monte Carlo simulations following an approach previously described by Blauch and Saveant to analyze redox polymers (20). The simulation approach (SI Appendix, *Materials and Methods*) randomly incorporates electron hopping and diffusion of the cytochromes on two-dimensional lattices with dimensions chosen to represent either the cylindrical surface of a whole cell or membrane extension. The key input parameters to each simulation are the time constant of electron hopping (*t*_*e*_), the time constant of physical motion (*t*_*p*_), and the fractional loading of cytochromes on the lattice (*X*, ratio of cytochrome density to maximum full packed density). The simulation output is the overall electron transport rate along the membrane surface.

The ratio *t*_*e*_*/t*_*p*_ plays a critical role in determining the overall electron transport behavior in the collision-exchange mechanism (16, 20). When physical motion is faster than electron hopping (*t*_*e*_*/t*_*p*_ > 1, illustrated in Movie S4), redox molecules redistribute on the lattice rapidly between successive electron hops, and the overall transport behavior can be well approximated with a mean-field model (16, 20). We previously applied this mean-field approach to assess electron transport along membrane extensions in this scenario (16). However, when physical motion is slower than electron hopping (*t*_*e*_*/t*_*p*_ < 1, illustrated in Movie S5), electron transport is in the percolation regime, where fast conduction requires high enough fractional loading to open up a conduction channel from an interconnected network of cytochromes spanning the entire lattice. In our system, *t*_*e*_ (the electron residence time in the decaheme cytochromes) can be estimated from previous measurements and molecular simulations to be in the 10^−5^ to 10^−6^ s range (13, 51– 53). To find *t*_*p*_, we use *D*_*phys*_ = 10^−2^ to 10^−1^ μm^2^/s, based on our *in vivo* diffusion coefficients (Fig. 4 and 6); the latter value is particularly observed in fast diffusing MtrC and OmcA on cells (Fig. 5B,D), and is also supported by *ex vivo* measurements of the MtrCAB transmembrane conduit on supported lipid bilayers, measured via fluorescence recovery after photobleaching to be approximately *D* = 10^−1^ μm^2^/s (54). Thus, *t*_*p*_ = 10^−3^ to 10^−4^ s (Eqn. 4, SI Appendix, *Materials and Methods*). Since for our system *t*_*e*_*/t*_*p*_ is generally < 1, the simplified mean-field approach previously applied (16) is no longer justified, and calculating the overall electron transport requires a stochastic simulation to account for the diffusion and electron hopping events of all redox carriers.

We report the simulation results as electron transport rate along the surface of a whole cell or membrane extension, as a function of cytochrome fractional loading (Fig. 8). Each curve depicts the simulation results for a particular combination of electron hopping constant and diffusion coefficient of cytochromes at the lower and upper limits of the realistic range described above (*t*_*e*_ = 10^−5^ to 10^−6^ s and *D*_*phys*_ = 10^−2^ to 10^−1^ μm^2^/s). For both simulation geometries and all combinations of hopping/diffusion coefficients, the electron transport rates exhibit a strong dependence, increasing by 3-4 orders of magnitude as a function of cytochrome fractional loading, as expected for transport in the percolation regime (20). At higher fractional loading (e.g. *X* > 0.7), the choice of diffusion coefficient does not impact the overall electron transport rate; in this limit, there is less room for physical diffusion and conduction is largely controlled by the electron hopping rate. Conversely, at lower fractional loading (e.g. *X* < 0.3), the electron transport rate is less sensitive to the electron hopping rate; in this limit, direct electron hopping events are less frequent in a landscape with sparsely distributed cytochromes, and conduction is controlled by the physical diffusion of cytochromes. It is interesting to consider these simulation results in light of previous experimental estimates of cytochrome concentrations in *S. oneidensis*. Ross *et al*. (15) estimated a total per cell MtrC and OmcA concentration of 100,000 proteins/cell, equivalent to a surface density of up to 30,000 proteins/μm^2^ for typical cell dimensions, e.g. 2 μm length and 0.5 μm diameter. This surface density, which is likely an upper limit since it assumes full localization of cytochrome to the outer membrane, translates to the upper range of fractional loading (*X* > 0.5, Eqns. 7-8). In this range of *X* = 0.5 to 1, our simulations (Fig. 8) reveal significant electron transport rates, in the 10^4^ to 10^5^ s^-1^ range for whole cell and membrane extension surfaces, depending on the exact choice of electron hopping constant (*t*_*e*_). We note that while these simulations account for the diffusion of the outer membrane electron transport proteins, other redox molecules (e.g. periplasmic cytochromes and small molecules such as flavins) may also contribute, leading to higher electron transport along membrane surfaces.

**Fig. 8.**
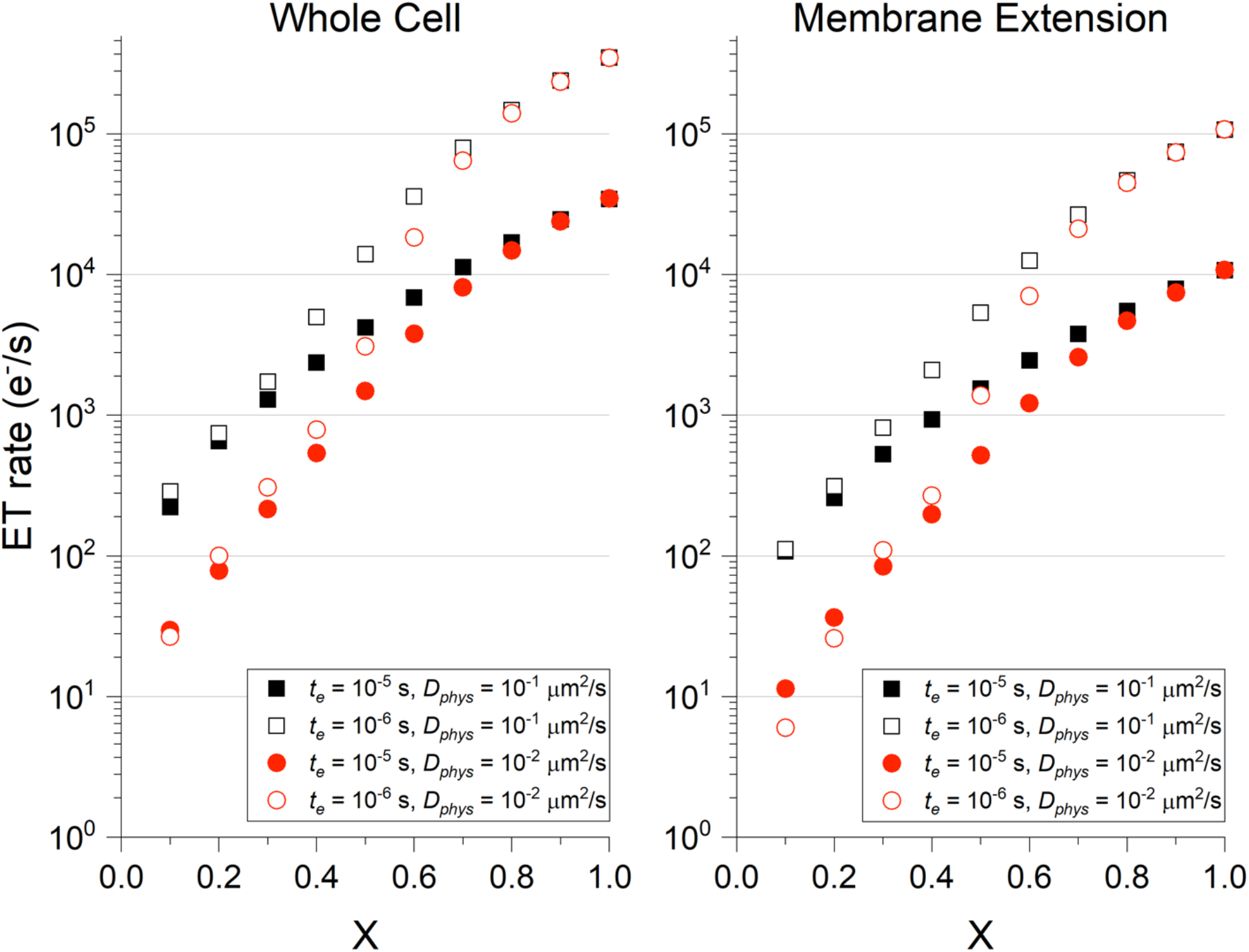
Simulation results of overall electron transport (ET) along the surface of whole cells or membrane extensions, based on experimentally measured diffusion coefficients. ET rates on the Y-axis are plotted on a log scale as a function of the fractional loading of redox carriers (*X*) on (*Left*) the surface of a whole cell (2 μm long and 0.5 μm in diameter) or (*Right*) the surface of an outer membrane extension (1 μm long and 100 nm in diameter). These results come from simulations using either *t*_*e*_ = 10^−5^ s (filled shapes) or 10^−6^ s (unfilled shapes) for a range of experimentally derived diffusion coefficients *D*_*phys*_ = 10^−1^ μm^2^/s (black squares) or 10^−2^ μm^2^/s (red circles).

To understand whether the simulated electron transport rates are consistent with experiments, we compare our results to existing measurements of the apparent electron diffusion coefficient (*D*_*ap*_) in electroactive biofilms and recent estimates of the redox conductivity (*σ*) in *S. oneidensis* biofilms. Using Fick’s law of diffusion, our calculated electron transport rate and the concentration gradient of reduced cytochromes along the cylindrical cell surface can be used to obtain *D*_*ap*_ (SI Appendix, *Materials and Methods*). Taking an electron transport rate of 10^4^ s^-1^ and a cytochrome concentration resulting from a representative fractional loading *X* = 0.5, this procedure results in *D*_*ap*_ ∼ 10^−8^ cm^2^/s, which is on the lower end of *D*_*ap*_ reported for electroactive bacterial biofilms (22). More recently, estimates of the redox conductivity of *S. oneidensis* have become available from electrochemical gating measurements of light patterned biofilms bridging interdigitated electrodes (55). From these measured conduction currents and using the full biofilm volume to define the conduction path (rather than only the cellular membrane surface), a biofilm conductivity (*σ*) of several nS/cm was estimated for *S. oneidensis*. Using the Nernst-Einstein relation (SI Appendix, *Materials and Methods*) to relate the apparent diffusion coefficient and conductivity (20), our calculated *D*_*ap*_ of approx. 10^−8^ cm^2^/s translates to *σ* ≈ 7 nS/cm, in remarkable agreement with the electrochemical gating measurements (55). These comparisons suggest that the simulated combination of electron hopping and cytochrome diffusion can explain many features of the observed redox conductivity of bacterial biofilms, at least in the case of *S. oneidensis*.

## Summary

In summary, we used single-particle tracking of quantum dot labelled multiheme cytochromes to demonstrate and quantify the lateral mobility of bacterial electron conduits on the cell surface and membrane extensions of *S. oneidensis* MR-1. The observed diffusive dynamics support a previously hypothesized role for cytochrome motion in facilitating long-distance electron conduction through collisions and electron exchange between the cytochromes along membrane surfaces. Based on these measurements, we performed kinetic Monte Carlo simulations that account for both electron hopping and physical diffusion of cytochromes. These simulations reveal significant electron conduction along cellular membranes and membrane extensions, with magnitudes that can explain experimental measurements of the apparent electron diffusion coefficient and electrical redox conductivity in bacterial biofilms. This study represents the first examination of the dynamics of bacterial electron conduits, and adds to a very limited data set on the diffusion of prokaryotic outer membrane proteins (23–25). The quantum dot labeling and tracking techniques demonstrated in this work can be used to study the importance and extent of diffusive dynamics of other bacterial cell surface proteins.

## Materials and Methods

### Strains, plasmids, and culture conditions

Bacterial strains and plasmids used or generated in this study are listed in Table 1. Generally, all Luria-Bertani (LB) agar plate cultures were grown overnight at 30°C for *S. oneidensis* or 37°C for *E. coli*, or up to 3 days at room temperature. All aerobic cultures were grown overnight in LB broth at 200 rpm and 30°C for *S. oneidensis* or 37°C for *E. coli*. All anaerobic *S. oneidensis* cultures were prepared by pelleting 5 mL of aerobic overnight LB pre-culture, washing in defined medium (17), and using it to inoculate 100 mL of anoxic defined medium in sealed serum bottles with 30 mM fumarate as the sole electron acceptor. These anaerobic cultures were then allowed to grow for approximately 24 hours at 30°C and 200 rpm where it reached late logarithmic phase (0.24-0.28 OD_600_). Frozen stocks of bacterial strains were stored in 30% glycerol at -80°C. Antibiotics (Kanamycin, 50 μg/mL) were added to media for bacterial cultures as needed to maintain selection of plasmid.

#### *In vivo* biotinylation

Anaerobically pre-grown *S. oneidensis* cells were harvested by centrifugation for 10 min at 7,142 × *g*, washed in PBS buffer supplemented with 5 mM MgCl_2_ (PBS-Mg) for 5 min at 4,226 × *g*, and resuspended in PBS-Mg, and collected in 1.5-mL tubes with 0.5 mL of cells diluted to 0.8 OD_600_ per sample. These samples were washed once again in PBS-Mg for 2 min at 7900 × *g*, and their supernatant was removed, leaving the cell pellet. The samples were then biotinylated *in vivo* using a BirA biotin-protein ligase standard reaction kit (Avidity). Following the kit instructions, each cell pellet was quickly resuspended in a 50-μL biotin ligase reaction mixture and left at room temperature for 1 h with vigorous shaking on an orbital shaker. Each 50-μL reaction mixture contained 50 mM bicine buffer (pH 8.3), 10 mM ATP, 10 mM MgCl_2_, 50 μM biotin, and 0.3 μM of BirA biotin ligase, dissolved in RNAse-free water. If necessary to prepare bigger samples, sample and reaction sizes were scaled up proportionately.

### Microscopy

Cells were prepared for *in vivo* microscopy by exogenous biotinylation as described above. Briefly, 0.5 mL of biotinylated cells initially diluted to 0.8 OD_600_ were washed 6 times in PBS at 12,000 rpm for 3 min each, resuspended in 0.5 mL of PBS and incubated for 1 h at room temperature under vigorous shaking with 50 μL of Qdot™ 705 Streptavidin Conjugate (SA-QD705, Thermo Fisher Scientific) prepared at 0.1-10 nM in PBS buffer + 6% BSA. Cells were then washed 3 more times in PBS and resuspended in 20 μL of PBS before mounting for microscopy.

Samples were mounted on high precision microscope glass coverslips (Marienfeld, #1.5, Ø25 mm) at the bottom of an open-air liquid imaging chamber. To promote cell attachment to coverslips, 5-10 μL of cells were dropped in the center of the coverslip and 1 mL of PBS was gently pipetted into the chamber. 5-10 min prior to imaging, FM 1-43FX membrane dye (Life Technologies; 0.0625-0.125 μg/mL) was added to the sample and gently pipetted to mix.

Imaging was performed on an inverted Nikon Eclipse Ti-E microscope equipped with total internal reflection optics, a 100× 1.49 NA objective (Nikon), two iXon Ultra EMCCD cameras (Andor Technology), a dual camera light path splitter (Andor Technology), and laser lines for excitation at 488 and 647 nm (Agilent). For splitting and simultaneously detecting FM 1-43FX and QD signals, a multiband pass ZET405/488/561/647x excitation filter (Chroma), a quad-band ZT 405/488/561/647 dichroic mirror (Chroma), and an emission splitting FF640-FDi01 dichroic mirror (Semrock) were used in combination with appropriate emission filters: ET525/50 (Chroma) for FM 1-43FX, and ET700/75 (Chroma) for QD705 and AF647. Channels were aligned prior to imaging using 40 nM TransFluoSphere streptavidin-labeled beads (488/645 nm, Life Technologies) as fiducial markers. Time-lapse microscopy was then performed in dual colors at an image acquisition rate of 40 ms/frame in each channel.

Generally, samples with 1-2 QDs per cell were imaged for tracking and diffusion analyses, as it facilitates signal localization and the building of trajectories over bacterial cells.

### Single-particle tracking

Single-particle localization and tracking were performed using SLIMfast, a program written for MATLAB that uses multiple-target tracing algorithms (56) and can accommodate for the blinking behavior of single molecules. The process of using SLIMfast for SPT has recently been described in detail for a study in *Caenorhabditis elegans* by (42). First, Fiji (ImageJ) software was used to convert the time-lapse microscopy data to TIFF image sequence files that could be opened by SLIMfast in MATLAB. Then, SLIMfast was used to localize single molecules based on 2D Gaussian fitting of the point spread function from each QD particle in each frame. Trajectories were then built by connecting the localized position of each particle over time from frame to frame, taking into account blinking statistics. Trajectories with at least 3 steps were used for diffusion analyses, as described in (42, 44) and in *SI Materials and Methods*. For tracking on the cell surface, at least 5,000-10,000 trajectories from 800-2,000 cells were typically analyzed for each condition; and only trajectories under 4 s in length were used for analysis, to further minimize the possibility of false connections between different quantum dots when building trajectories. For added quality checking, several fields in each cell dataset and all videos from each OME dataset were manually inspected frame-by-frame in Fiji (ImageJ) to verify overlay of particle localizations (specifically those localizations used to generate trajectories, as some localizations are discarded during the tracking process) with raw signal from QDs. Similarly, those fields were also inspected to ensure that localizations used for trajectories were found within cells, as labeled by raw signal from the membrane dye. Also, if needed to remove stray trajectories from clearly moving cells, or to limit the region of interest to an OME, regions of interest (ROI) were defined in SLIMfast, and localization and tracking were repeated with those ROI to obtain final trajectories for analysis.

## Data availability

MATLAB scripts written for this paper are available to readers and are shared publicly on the Open Science Framework repository (https://osf.io/c49uw/) (59).

## Acknowledgements

We thank Dr. Jeffrey A. Gralnick (University of Minnesota) whose lab sent us the *S. oneidensis* MR-1 Δ*mtrC* and Δ*omcA* deletion mutants, as well as the pBBR1-MCS2 plasmid (in *E. coli* host). We thank Dr. Thomas A. Clarke (University of East Anglia) who advised us on positioning the AP tag. We thank Dr. Namita P. Schroff and Dr. Steven E. Finkel (University of Southern California) who gave early advice on molecular biology techniques, and Dr. James Boedicker’s lab (University of Southern California) for providing molecular biology resources. Lastly, thanks to Magdalene MacLean, who assisted G.W.C. with various tasks.

Measurements of the cytochrome diffusive dynamics in M.E.-N’s laboratory were supported by the Division of Chemical Sciences, Geosciences, and Biosciences, Office of Basic Energy Sciences of the U.S. Department of Energy through grant DE-FG02-13ER16415. G.W.C. also acknowledges support by the National Science Foundation Graduate Research Fellowship Program (grant DGE1418060). Y.Z. was partially supported by the NIH T32 Chemistry Biology Interface Training Grant at the University of Southern California. S.P. was initially supported by Air Force Office of Scientific Research award FA955014-1-029 and then by the US Office of Naval Research Multidisciplinary University Research Initiative Grant No. N00014-18-1-2632.

## Author contributions

M.Y.E-N and F.P. conceived the project. G.W.C. designed, performed and analyzed experiments, with advice from M.Y.E-N. and all other authors (S.P., Y.Z., L.A.Z. and F.P.) based on area of expertise. S.P. and M.Y.E-N. shared physics/math expertise. L.A.Z., M.Y.E-N. and F.P. shared biology expertise. Y.Z. and F.P. provided tools and assistance with microscopy, tracking, and diffusion analyses. S.P. designed, performed, and analyzed all simulations. G.W.C. and M.Y.E-N. wrote and edited the paper, with feedback from S.P., Y.Z., L.A.Z., and F.P.

The authors declare no conflict of interest.

