## Supplementary Material for "Single molecule tracking of bacterial cell surface cytochromes reveals dynamics that impact long-distance electron transport"

**This file includes:**

SI Materials and Methods
Tables S1 to S6
Figs. S1 to S7
Captions for Movies S1 to S5

**Other Supporting Information for this manuscript includes the following:**

Movies S1 to S5

### SI Materials and Methods

#### Plasmid construction

Genetic constructs were generated to add a 15-amino acid biotin acceptor peptide (AP) tag sequence (1–4) to the C-termini of outer membrane-associated cytochromes MtrC and OmcA in *S. oneidensis* MR-1. Plasmid design is illustrated in SI Appendix, Fig. S1 and S2 and their construction is described here.

To make DNA inserts encoding the AP-tagged genes, *S. oneidensis* MR-1 genomic DNA template was first obtained from stationary phase LB cultures using a Mo Bio UltraClean Microbial DNA Isolation Kit. Primers listed in SI Appendix, Table S1 were used in three consecutive rounds of overhang extension PCR to add the AP tag (DNA sequence CTCGTGCCACTCGATCTTCTGGGCCTCGAAGATATCGTTCAGGCC; amino acid sequence GLNDIFEAQKIEWHE) connected by a serine-glycine linker (DNA sequence TGAACC, with serine closer to the AP tag), to the end of each gene immediately before the stop codon. Forward primers were also designed to amplify the native promoter region of each gene (118 bp upstream of *mtrC* or 114 bp upstream of *omcA*, respectively). Primers were also designed to include restriction sites for XhoI and XbaI, plus 5-bp of protection bases at outside ends of each DNA insert. All primers were synthesized by Integrated DNA Technologies and obtained as desalted oligonucleotides. All PCR was performed using a Bio-Rad C1000 Touch™ Thermal Cycler, and protocols for each round of overhang PCR are listed in SI Appendix, Tables S2-S4 and are based on manufacturer's instructions for Phusion High Fidelity DNA polymerase (New England Biolabs). Annealing temperatures were calculated by the New England Biolabs annealing temperature (Tm) calculator (http://tmcalculator.neb.com/#!/main). For each 35-cycle PCR run, the first 5 cycles used the annealing temperature calculated for only the template-binding portions of each primer pair, and the last 30 cycles used the annealing temperature calculated for the full-length primer pairs. After each PCR experiment, a portion of the sample was separated on a DNA gel (1% agarose in TBE buffer, run for ~60 min at 100 V) to check if the PCR worked (i.e. amplified DNA of appropriate size). All DNA gels were run on a Bio-Rad Mini-Sub Cell GT electrophoresis system followed by visualization on a Bio-Rad Gel Doc EZ™ Imager. If PCR was successful, then the PCR product was purified using a Purelink® PCR Purification Kit for use as template for the next round of extensions via overhang PCR. Finally, after generating the full-length DNA inserts by PCR, they were purified using QIAquick Gel Extraction Kit (Qiagen). All PCR reactions were 25 µL in volume, except the final extension was repeated with 100 µL in reaction volume to make more DNA insert for gel purification. Generally, all genomic DNA was stored at 4°C and all primers and PCR products were stored at -20°C in between steps. All DNA concentrations were measured using a Thermo Scientific Nanodrop 2000c spectrophotometer.

The DNA inserts were cloned via restriction digest into the pBBR1-MCS2 broad host cloning vector (5), which is Kanamycin resistant and can also be used for blue-white screening (aka β-galactosidase assay) in *E. coli*. Plasmid samples were purified from overnight LB cultures of its *E. coli* host using Purelink Quick Plasmid Miniprep Kit (Invitrogen). Restriction digestion reactions using XhoI and XbaI (New England Biolabs) were performed according to manufacturer's guidelines. Then, the digested plasmid and insert were immediately combined in the recommended 1:3 vector:insert ratio for ligase reactions, following manufacturer's protocol (New England Biolabs Quick Ligation Kit #M2200). Ligation reaction volumes were scaled up to however much digested insert/plasmid DNA was available. Finally, the freshly ligated constructs were plasmid purified and stored at 4°C before transformation via electroporation.

#### Transformation and quality checking

First, electrocompetent *E. coli* DH5α cells were freshly prepared on ice. For each sample, 1 mL of an overnight LB culture was centrifuged at 7900 × *g* for 1 min, washed 3x by gentle pipetting in chilled (4°C) 10% glycerol at 7900 × *g* for 2 min. The last 50-70 µL of liquid from the 3rd wash was left in each sample for final resuspension. The electrocompetent cells were then incubated with varying amounts (1-10 µL) of purified, ligated DNA (approx. 5-60 ng in total) for 2 min on ice. Samples were electroporated in an Eppendorf Eporator® machine at 1.7-2 kV with a time constant of approx. 5 ms, quickly resuspended in 0.5-1 mL of fresh LB broth, and allowed to recover in a 37°C shaking incubator for 90 min. Afterwards, various amounts (50-200 µL) of sample were spread onto LB agar supplemented with Kanamycin for antibiotic selection and X-gal (5-bromo-4-chloro-3-indolyl-β-D-galactopyranoside, 20 mg/mL in dimethylformamide, 60 µL spread on agar plates) for blue-white screening (aka β-galactosidase assay). If colonies grew and appeared white on LB + Kan + X-gal plates, that suggested they had received plasmid containing our DNA insert, so up to 20 apparently white colonies were re-streaked onto new LB + Kan + X-gal plates 2-3 times to make sure colonies consistently appeared white rather than pale or dark blue. As a control, a no-DNA sample was also included in each electroporation experiment and plated on both LB agar + Kan (no growth expected) and LB agar (growth expected).

Colonies that consistently appeared white during blue-white screening (i.e. possibly successful transformants) were then used for PCR and DNA gel electrophoresis. Here, we used colony PCR (using appropriate Forward and Reverse 1 Primers from SI Appendix, Table S1) to see if we could amplify respective DNA inserts from the white transformant colonies. To do colony PCR, a sterile pipet tip was used to pick up a portion of a colony and vigorously resuspend it in 10 µL of RNAse-free water. Then, 1 µL of this mixture was used as template for PCR. For positive controls, 1 µL of purified DNA insert (generated by overhang extension PCR and stored in -20°C from previous experiments) was used as template for PCR and used as a comparison for desired DNA size in DNA gels. Detailed protocols for colony PCR are listed in SI Appendix, Tables S5-S6 and are based on manufacturer's instructions for OneTaq Quick-Load DNA Polymerase (New England Biolabs). For each 30-cycle PCR run, annealing temperatures for the first 5 cycles were calculated for only the template-binding portion of each primer, while the last 25 cycles were calculated for the full-length PCR primers. Based on DNA gels of the PCR products, samples that clearly amplified the desired DNA insert were cultured overnight in 5 mL LB broth and stored in 20% glycerol at -80°C to be used in further experiments.

Plasmids (encoding putative AP-tagged MtrC or OmcA) purified from these *E. coli* transformant cultures were then used for transformation into *S. oneidensis* MR-1 in their respective gene deletion backgrounds Δ*mtrC* or Δ*omcA* (6) using a recently developed electroporation protocol (7). Control samples with no DNA were also included in each experiment and plated on both LB agar + Kan (no growth expected) and LB agar (growth expected). Several transformants which grew on LB agar + Kan were then grown overnight in LB Broth + Kan. Then, similarly to quality checking the *E. coli* transformants, the *S. oneidensis* transfomants were checked by PCR (using appropriate Forward and Reverse 1 Primers from SI Appendix, Table S1) and DNA gel electrophoresis to see if we could amplify respective DNA inserts from the transformants. As before, samples using the desired DNA insert were used as a positive control. Finally, transformants verified by PCR were frozen at -20% glycerol and stored at -80°C to be further verified by DNA sequencing and SDS-PAGE heme staining.

#### DNA sequencing

Sanger sequencing was performed to verify if successful transformants contained the correct sequence for the desired AP-tagged genes. To prepare samples for sequencing, we amplified the region containing the DNA inserts from purified plasmid DNA from overnight LB cultures. Here, we used standard M13 Forward and Reverse primers (Integrated DNA Technologies, also listed in SI Appendix, Table S1) and performed PCR according to manufacturer's instructions as described for Phusion High Fidelity DNA Polymerase (New England Biolabs). Purified PCR products were then mixed with M13 Forward primer and sent to GeneWiz for sequencing. Sequencing results were subsequently viewed by SnapGene Viewer software.

#### SDS-PAGE and heme staining

To prepare lysed cell samples for SDS-PAGE gel electrophoresis, LB cultures were aerobically grown overnight, centrifuged as 1 mL samples at 7900 × *g* for 2 min, supernatant discarded, and resuspended in 100 µL of sample buffer with reductant and 25 µL of 4x loading dye. To make stock sample buffer, we mixed 3.55 mL distilled water, 1.25 mL 0.5 M Tris HCl, pH 6.8, 2.5 mL glycerol, 2.0 mL 10% (w/v) SDS, and 0.2 mL 0.5% (w/v) bromophenol blue. Sample buffer with reductant was made fresh before each experiment adding 50 µL of β-mercaptoethanol to 950 µL of stock sample buffer. Then samples were boiled in a hot water bath for 5-10 min at 95-100°C before storage at -20°C or immediate use in gel electrophoresis. Samples were loaded onto SDS-PAGE gels (previously prepared using Bio-Rad TGX FastCast Acrylamide Kit) and run at 200 V for ~1 h on a Bio-Rad Mini-PROTEAN Tetra Cell System. Our electrophoresis running buffer contained 3.03 g/L Tris base, 14.4 g/L glycine, and 1 g/L SDS, pH 8.3. Samples were typically run on replicate gels. One gel was reserved for staining with Instant Blue™ (Expedion) total protein dye. The other gels were then visualized with peroxidase activity-based heme staining techniques.

Heme staining via peroxidase activity assay with 3,3'-diaminobenzidine (DAB) was adapted from a previously described procedure (8). After electrophoresis, gels were immediately pre-incubated for 30 min at room temperature in a solution containing freshly made 25 mg DAB in 50 mL of 0.5

M Tris Buffer, pH 7 (sonicated as needed up to 1 h). Then, this solution was discarded, and the gels were incubated 30 min to overnight at 4°C in a second solution containing peroxide (50 mg DAB in 50 mL of 50 mM Tris HCl, pH 8, sonicated as needed up to 1 h, with 1.5 mL of 30% hydrogen peroxide added right before use). Staining was generally visible within 30 min after adding peroxide.

#### Western blot

Western blot experiments were also performed to test that *in vivo* exogenous biotinylation worked and that this labeling was specific to tagged proteins of interest. Freshly biotinylated cell samples were washed 2x in PBS buffer and prepared for SDS-PAGE gel electrophoresis as described above. While SDS-PAGE gels were running, fiber pads, filter paper, and nitrocellulose membrane were cut to appropriate gel size and soaked for >20 min in chilled transfer buffer containing 25 mM Tris, pH 8.3, 192 mM glycine, and 20% v/v methanol. After SDS-PAGE, gels were also soaked in chilled transfer buffer for >20 min. Then, proteins were transferred from gel to membrane in a Bio-Rad Mini Trans Blot Transfer Cell system run at 100 V and 350 mA for 1 h, in chilled and stirred transfer buffer. Afterwards, the membrane was removed and incubated in 10 mL of blocking solution made from 3% BSA in TBST (20 mM Tris-HCl, 140 mM NaCl, pH 7.5, 0.1% v/v Tween-20) and left for 1 h at room temperature or overnight at 4°C. After blocking, the membrane was washed 3x in TBST for 5 min with gentle shaking. Then, it was incubated with diluted polyclonal rabbit-raised MtrC or OmcA-specific primary antibody (9) (rαMtrC: 0.66 µg/mL, rαOmcA: 0.49 µg/mL, both in 1% BSA/TBST) at room temperature for 30 min with gentle shaking. After 3 more 5-min washes in TBST, the membrane was then incubated in horseradish peroxidase (HRP)-conjugated goat anti-rabbit secondary antibody (Abclonal) diluted 1:2000 in 1% BSA/TBST for 30 min at room temperature with gentle shaking. Since the HRP is light sensitive, this incubation and future steps were done in a covered container. Then the membrane was washed 3x in TBST, 3x in TBS, and 1x in water for 5 min each with gentle shaking. Finally, the membrane was visualized using SuperSignal™ West Pico PLUS Chemiluminescent Substrate (varying incubation time and exposure time as appropriate) and developed on X-ray film. Then, we prepared the blots for restaining by using a mild stripping procedure adapted from Abcam: blots were incubated 2x for 5-10 min in mild stripping buffer (For 1 L: 15 g glycine, 1 g SDS, water, adjusted to pH 2.2) at room temperature over gentle shaking, then washed 2x for 10 min in PBS, and 2x for 5 min in TBST. The stripped membranes were then blocked for 1 h in 3% BSA/TBST and washed 3x for 5 min in TBST. Then, they were incubated in HRP-conjugated streptavidin (Thermo Fisher Scientific) diluted to 0.25 µg/mL in 1% BSA/TBST, gently shaking at room temperature for 30 min. Finally, the membranes were washed 3x in TBST, 3x in TBS, and 1x in water for 5 min each and visualized with chemiluminescent substrate and developed on X-ray film as described above.

The first set of films were used to detect proteins of interest using MtrC and OmcA-specific antibodies (though there is some nonspecific binding, where MtrC antibody also binds to OmcA, and vice versa). Since the streptavidin binding is not reversible, this was probed last. The second set of films, which captured signal from streptavidin, revealed which proteins are biotinylated, and could be compared to the first set to verify overlap of signal, and thus specificity of biotinylation to proteins of interest.

#### Microscopy

For initial labeling control experiments (Fig. 2B and SI Appendix, Fig. S4B), 20 nM streptavidin-conjugated Alexa Fluor 647 (Thermo Fisher Scientific) was used instead of SA-QD705. Other steps for sample preparation and microscopy were as described under *Materials and Methods*.

For the cell-free QD experiment (SI Appendix, Fig. S5), SA-QD705 was diluted to a concentration of 0.002 nM in PBS buffer, or sufficient dilution to resolve individual QDs in microscopy. Then, 0.5 mL of this sample was mounted on a glass coverslip, and a total of 10 microscopy fields (for a total of >1000 individual QDs) were imaged. Other imaging and tracking steps were performed as described for other experiments under *Materials and Methods*.

For the biotin flood experiment (SI Appendix, Fig. S6), the quantum dot labeling step was modified to "flood" the sample with excess biotin shortly after addition of SA-QD705. The purpose of this step was to saturate streptavidin binding sites on the QD probes and prevent individual probes from binding to multiple protein targets. Briefly, 5 min after addition of SA-QD705, excess biotin (≥200-fold biotin to QD ratio) dissolved in PBS buffer was added. The required amount of excess biotin was calculated by the following estimations: assuming 5-10 wild type streptavidin molecules per QD based on manufacturer estimates, 10 streptavidin molecules would have 40 biotin sites, requiring at least a 40-fold ratio of excess biotin to QD, in order to saturate remaining streptavidin-biotin binding sites. To obtain excess biotin at a ≥200-fold biotin to QD ratio, we added 200 µL of 1 µM biotin to a 50-µL sample containing 20 nM QD. Samples were incubated at room temperature with shaking for 15 min prior to final wash steps and imaging. Other steps for sample preparation, imaging, and tracking were performed as described under *Materials and Methods*.

#### Diffusion analyses

As mentioned in *Materials and Methods*, single-particle localization and tracking was performed using SLIMfast, a program written for MATLAB that uses multiple-target tracing algorithms (10) and can accommodate for the blinking behavior of single molecules. SLIMfast was also used to extract data for diffusion analyses, as recently described for a study in *C. elegans* (11).

Probability distribution of square displacement (PDSD) analyses were performed in SLIMfast on all trajectories to determine the number of major populations of diffusing behavior for each dataset. Briefly, cumulative probability distribution functions of square displacements $P\ (r^2)$ for selected $\Delta t$ were generated in SLIMfast and fitted with a model for $i$ = 1, 2, or 3 (or more) components (aka populations or diffusion behaviors) within the overall dataset. The cumulative probability curve defines the probability $P$ that a particle is displaced by a distance $r$ from its original position given a certain period of time $\Delta t$. The fitting model is described in detail by

Schütz *et al.* under their equations 3 (model for 1 component) and 5 (model expanded for 2 components) and can be similarly expanded for 3 or more components (12). The general model based on Schütz *et al.* (12) is described below and in (11, 13, 14):

$$P(r^2, \Delta t) = 1 - \sum_{i=1}^{n} \alpha_i e^{-r^2/r_i{}^2(\Delta t)}$$

$$\sum_{i=1}^{n} \alpha_i\,(\Delta t) = 1 \qquad \text{(Eqn. 1)}$$

Here, fitting coefficients $\alpha_i(t)$ and $r_i{}^2(t)$ are functions of time lag $\Delta t$, where $\alpha_i(\Delta t)$ is the fraction of square displacements $r_i{}^2(\Delta t)$ corresponding to a particular component $i$ at a given value of $\Delta t$. When performed in SLIMfast, this analysis is also used to sort the raw displacement data into populations which can then be extracted for ensemble MSD curve analysis, described below.

Ensemble MSD curves were then generated for each diffusing population in each dataset (extracted by PDSD analysis in SLIMfast). Ensemble MSD curves are also called time-averaged MSD curves or $r^2(\Delta t)$ curves (mean squared displacement $r^2$ as a function of time lag $\Delta t$). The shape of this curve indicates the overall diffusing behavior (e.g. free, confined) of that population, where free (Brownian) diffusion is characterized by a straight upward slope, and confined diffusion is characterized by a slope that bends to a plateau over time. Subsequently, the curve can be fitted with appropriate diffusion model (e.g. free, confined) which yields information of interest such as diffusion coefficient *D* and confinement radius *R*. SLIMfast was used to calculate ensemble MSDs from raw displacement data and the MSD curves were then plotted in Origin 2019b software. Error bars at each time point of the ensemble MSD curve show Y-error = $\pm\frac{r^2}{\sqrt{N}}$, where *N* is the number of independent data points (i.e. displacements) that were analyzed to give the mean squared displacement $r^2$ value for each time interval $\Delta t$.

If appropriate, diffusion coefficients (*D*) for confined diffusion were determined by fitting ensemble MSD curves with a circularly confined diffusion model:

$$r^2 = R^2\left(1 - A_1 e^{\frac{-4A_2 D\Delta t}{R^2}}\right) + 4\sigma^2 \qquad \text{(Eqn. 2)}$$

Here, $r^2$ is the square displacement (or MSD), *D* is the diffusion coefficient, $\Delta t$ is the time lag or time interval, *R* is the corral/confinement radius, $\sigma$ is the position error, and $A_1$ = 0.99 and $A_2$ = 0.85 (constants defined by corral geometry, in this case circular confinement), as described in (13, 15).

If appropriate, diffusion coefficients for free (Brownian) diffusion were determined by fitting ensemble MSD curves with the following model, accounting for position error:

$$r^2 = 4D\Delta t + 4\sigma^2 \quad \text{(Eqn. 3)}$$

To build histogram distributions of diffusion coefficients, individual MSD curves for individual trajectories were generated in SLIMfast. Individual diffusion coefficients were calculated by fitting each MSD curve for the first 3 time lags of the trajectory with the model for free diffusion (Eqn. 3). Diffusion coefficients were then assembled in histograms using Origin 2019b software.

All diffusion coefficients are reported in micrometer squared per second ± standard deviation (SD) of the fit value. Significant differences between diffusion coefficients were determined by F-tests.

#### Kinetic Monte Carlo simulations of long-distance electron transport

Simulations of the overall electron transport (ET) along the cell surface or membrane extensions are dependent on the relative rates of direct electron hopping vs. redox carrier diffusion, characterized by the ratio $t_e/t_p$, where $t_e$ and $t_p$ are the time constants of electron hopping and physical motion of redox carriers, respectively (16). These time constants are related to the diffusion coefficients for electron hopping between redox carriers ($D_e$), and the physical motion of the redox carriers ($D_{phys}$) according to the following equations:

$$D_e = \frac{\delta^2}{4t_e}, D_{phys} = \frac{\delta^2}{4t_p} \quad \text{(Eqn. 4)}$$

where $\delta$ is the center-to-center distance of closest approach between redox carriers and is estimated as the average size of one redox carrier.

For our system, $t_e$ is estimated by the electron residence time in outer membrane decaheme cytochromes to be approximately $10^{-5}$ to $10^{-6}$ s from molecular simulations of the electron flux (17–19), and $t_p$ can be estimated from our diffusion measurements of MtrC and OmcA in this study using Eqn. 4; taking $\delta$ = 6.33 nm for MtrC (20), $D_{phys}$ values of $10^{-2}$ to $10^{-1}$ µm$^2$/s translate to a $t_p$ of $10^{-3}$ to $10^{-4}$ s.

Since $t_e < t_p$ in our system, we ruled out the mean-field approach discussed in (16, 21), and instead performed kinetic Monte Carlo simulations in MATLAB that randomly simulate direct electron hopping and redox carrier diffusion, following the Blauch–Saveant approach (16) for modeling electron transport in an assembly of redox carriers in two dimensions. These simulations are illustrated in Movies S4 and S5. In these simulations, the cylindrical surface of a cell or membrane extension with length $L$ and diameter $d$ was represented by a 2D lattice array, where one dimension of the lattice ($y$-axis) was the length ($L$) of the surface (cell or membrane extension) and the other ($x$-axis) was its circumference ($\pi d$). To account for a representative cell surface we used $L$ = 2 µm, $d$ = 0.5 µm. To account for a representative outer membrane extension, we used $L$ = 1 µm, $d$ = 100 nm. The size of a single lattice site (i.e., a possible redox carrier position) was estimated as $\delta^2$, i.e. a square based on the average diameter of a redox carrier. The total number of sites in the lattice was chosen by dividing its surface area ($L\pi d$) by the surface area of a single lattice site ($\delta^2$).

At the beginning of each simulation, this 2D array was populated by redox carriers that were randomly distributed on the surface at a given fractional loading $X$. The reduced/oxidized state of the redox carriers was initially set up to give a linear gradient of reduced redox carriers along the main axis ($y$-axis) to decrease the simulation time needed to reach steady state. In each simulation time step, the redox carriers were free to diffuse randomly in any direction (left, right, up, or down), from a current occupied position to an adjacent unoccupied position. Similarly, electrons could hop in any direction (left, right, up, down), from a reduced redox carrier to an adjacent oxidized redox carrier. The left and right side edges of the lattice were connected through periodic boundary conditions to allow unrestricted movement along the $x$-axis (i.e., circumference). In each time step, redox carriers were reduced at one end of the main axis ($y$-axis) and oxidized at the other end. The number of oxidation events (i.e., number of electrons transferred) at the end of the lattice was counted over time. This number (i.e. the net number of electrons that crossed the length of the lattice per unit time) is defined as the overall electron transport rate ($I$) and is dependent on the width of the given 2D lattice.

To achieve a high enough temporal resolution for accurate simulation of hopping and diffusion events, the simulation time step must be much smaller than the smallest time constant ($t_e$ or $t_p$). Since in our case $t_e$ was the smaller time constant, the time step ($\Delta t$) was chosen to be $\Delta t = 0.1 \times t_e$ in all simulations. Each simulation was run long enough to achieve steady state, leading to a constant electron transport rate ($I$) (in electrons per second) which was used as the simulation output. The number of steps used in our simulations ranged from $10^6$ to $4\times10^7$. The most important parameter affecting the number of required steps was $t_p$ (and thus $D_{phys}$), with larger $t_p$ values (smaller $D_{phys}$) requiring a longer simulation.

Using Eqn. 5, we can then normalize $I$ by the width of the 2D lattice (i.e., the circumference of the cell or membrane extension) to find the electron flux ($J$):

$$J = \frac{I}{\pi d} \quad \text{(Eqn. 5)}$$

If desired, $J$ can then be used to find $D_{ap}$ using Fick's first law of diffusion:

$$J = D_{ap}\frac{\partial C}{\partial x} \approx D_{ap}\frac{C(x=0) - C(x=L)}{L} = D_{ap}\frac{C_0}{L} \quad \text{(Eqn. 6)}$$

where $C$ is the concentration of reduced redox carriers, $L$ is the length of the lattice along the main axis ($y$-axis), and $C_0$ is the concentration of all redox carriers (16, 21). $C_0$ can be calculated for a given fractional loading $X$, based on the definition of fractional loading:

$$X = \frac{C_0}{C_{Max}} \quad \text{(Eqn. 7)}$$

where $C_{Max}$ is the maximum possible concentration of redox carriers on a membrane surface. Assuming spherical redox carriers, $C_{Max}$ can be estimated by the concentration of circles in a hexagonal lattice, $C_{Hex}$:

$$C_{Max} \approx C_{Hex} = \frac{0.91}{\pi\left(\frac{\delta}{2}\right)^2} \quad \text{(Eqn. 8)}$$

Eqns. 7-8 can be used to find $C_0$ for a given input value of $X$. Then, we can rearrange Eqn. 5-6 to calculate a general $D_{ap}$ for our system:

$$D_{ap} = \frac{IL}{\pi d C_0} \quad \text{(Eqn. 9)}$$

If desired, the Nernst-Einstein relation can be used to relate $D_{ap}$ to conductivity ($\sigma$), as discussed by (16):

$$\sigma = \frac{D_{ap} C_0 e^2}{kT} \quad \text{(Eqn. 10)}$$

Here, $e$ is the charge of 1 electron, $k$ is Boltzmann's constant, and $T$ is the temperature. $C_0$ refers to the number of charge carriers (i.e., the concentration of redox carriers) which we now estimate for a 3D volume (cylinder volume: $L\pi\left(\frac{d}{2}\right)^2$) rather than a 2D lattice area (cylinder surface area: $L\pi d$), using $C_0 = XC_{max}\frac{4}{d}$, where $d$ is the diameter of a cell (typically 0.5 µm).

#### SI Figures and Tables

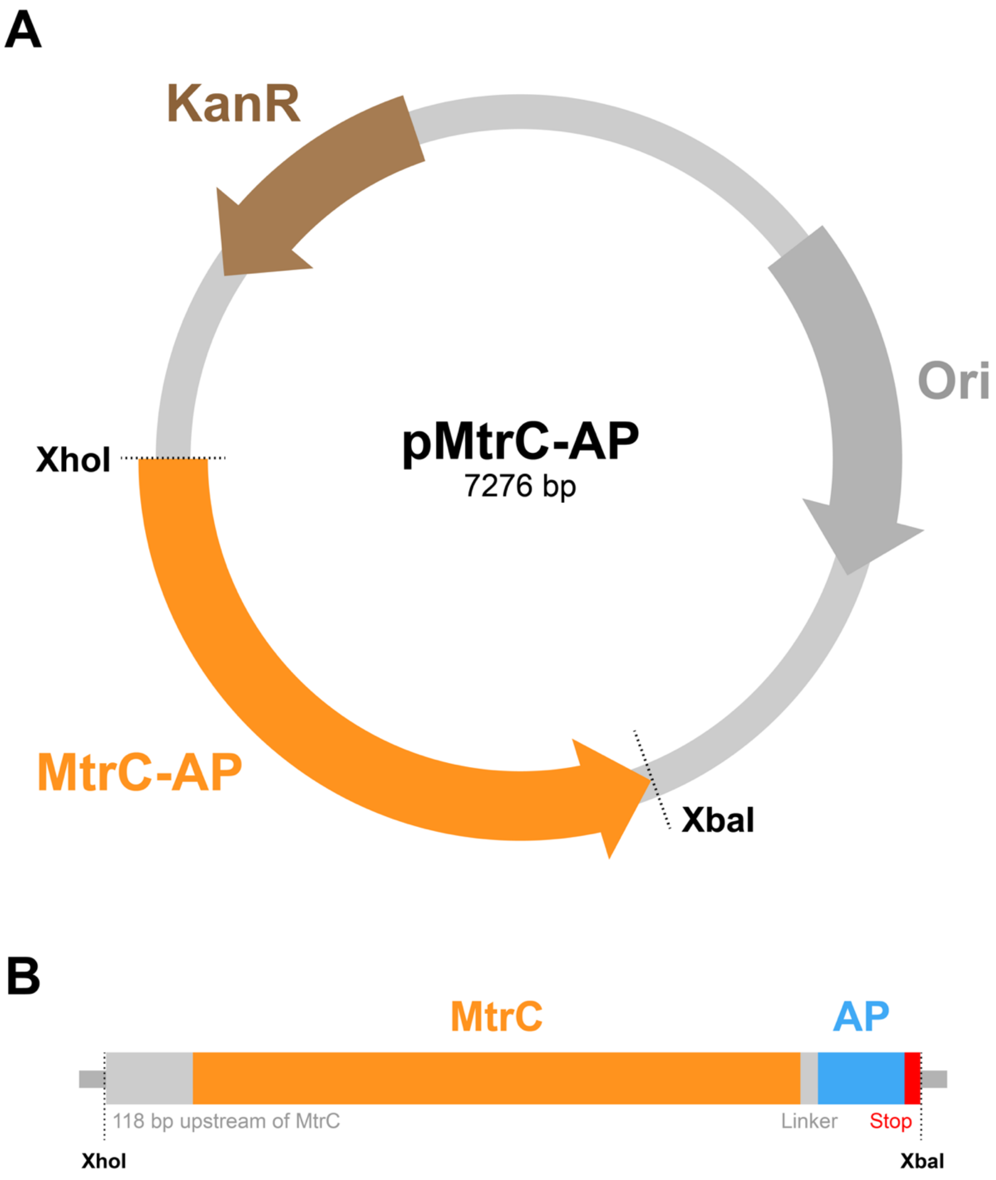


**Fig. S1. Schematic of design for MtrC-AP construct, dubbed pMtrC-AP. (*A*)** Plasmid design. MtrC-AP gene fusion was inserted between restriction sites for XhoI and XbaI in pBBR1-MCS2 plasmid (5) with kanamycin resistance. **(*B*)** Insert design. Here, the C-terminus of MtrC was fused to a 45-bp "AP Tag" encoding the 15-amino acid biotin acceptor peptide (AP: GLNDIFEAQKIEWHE) from *E. coli*, as described in (3, 4). Insert was generated from *S. oneidensis* MR-1 genomic DNA template using three rounds of overhang PCR with primers for MtrC listed in SI Appendix, Table S1. DNA insert included an XhoI restriction site, 118 bp upstream of *mtrC* (including native promoter), protein-coding region for MtrC, a short glycine-serine linker, and the 45-bp biotin acceptor peptide (AP) tag sequence just before the stop codon and XbaI restriction site. Total length of DNA inserted into pBBR1-MCS2 plasmid was 2,185 bp for a final construct size of 7,276 bp.

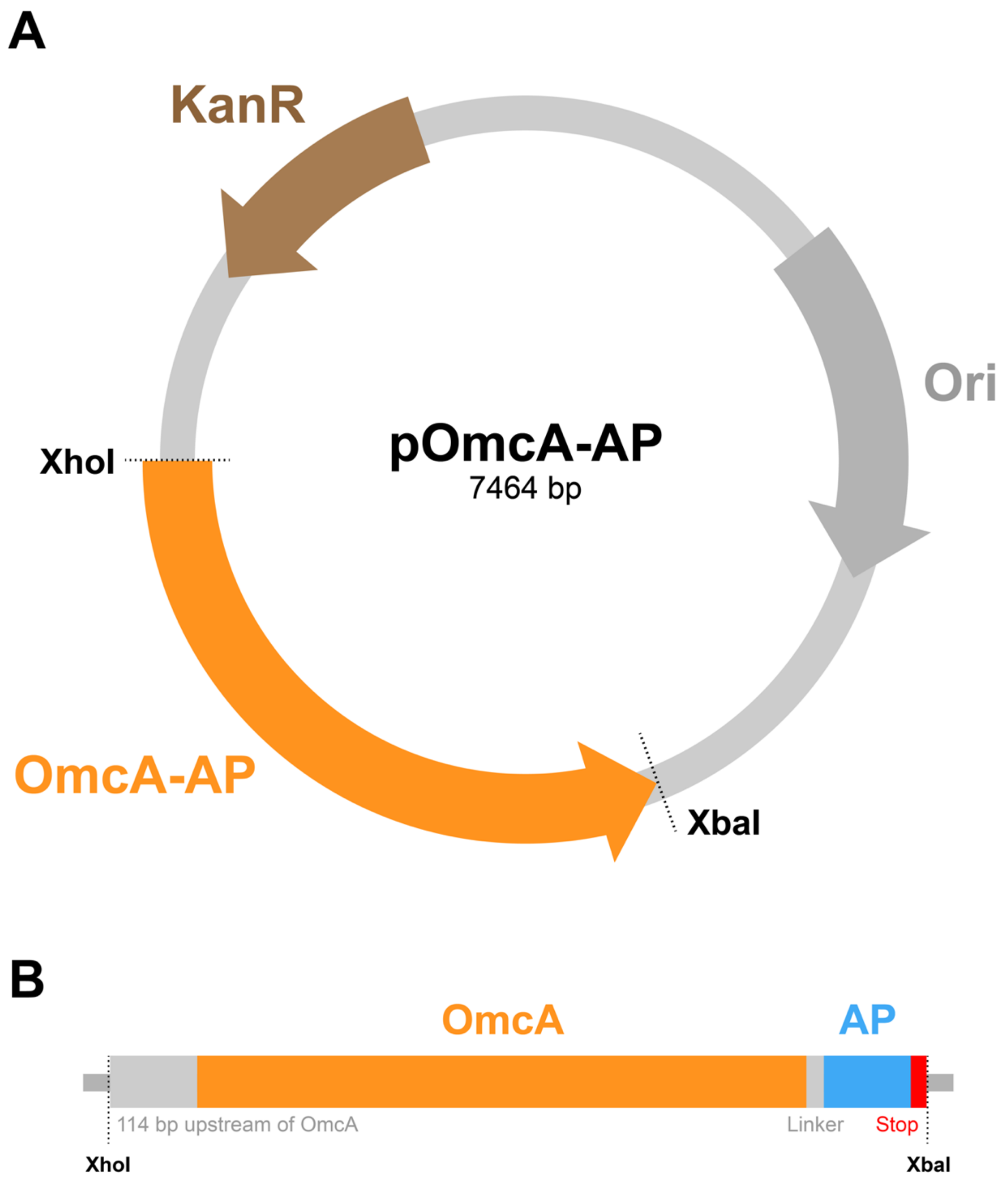


**Fig. S2. Schematic of design for OmcA-AP construct, dubbed pOmcA-AP. (*A*)** Plasmid design. OmcA-AP gene fusion was inserted between restriction sites for XhoI and XbaI in pBBR1-MCS2 plasmid (5) with kanamycin resistance. **(*B*)** Insert design. Here, the C-terminus of OmcA was fused to a 45-bp "AP Tag" encoding the 15-amino acid biotin acceptor peptide (AP: GLNDIFEAQKIEWHE) from *E. coli*, as described in (3, 4). Insert was generated from *S. oneidensis* MR-1 genomic DNA template using three rounds of overhang PCR with primers for OmcA listed in SI Appendix, Table S1. DNA insert included an XhoI restriction site, 114 bp upstream of *omcA* (including native promoter), protein-coding region for OmcA, a short glycine-serine linker, and the 45-bp biotin acceptor peptide (AP) tag sequence just before the stop codon and XbaI restriction site. Total length of DNA inserted into pBBR1-MCS2 plasmid was 2,373 bp for a final construct size of 7,464 bp.

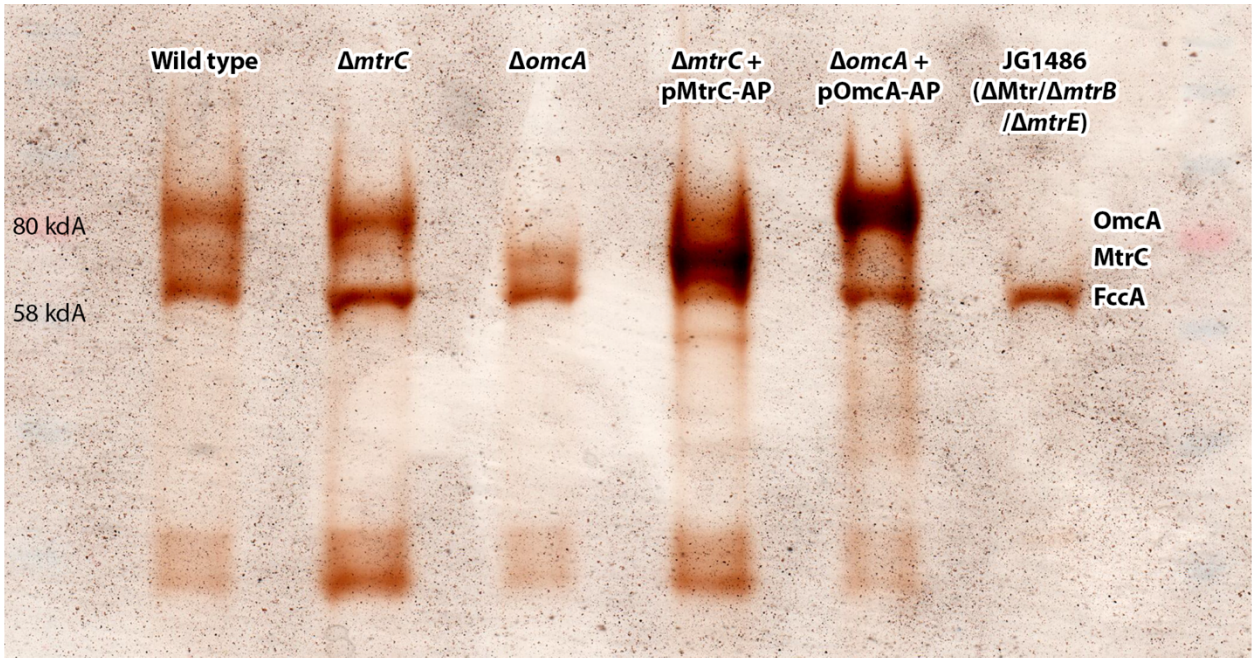


**Fig S3. Heme staining of protein gels via peroxidase activity assay reveals heme in newly tagged cytochromes MtrC-AP and OmcA-AP**. Redox-dependent staining using a 3,3'-diaminobenzidine (DAB) and hydrogen peroxide ($H_2O_2$) peroxidase activity assay. SDS-PAGE and subsequent staining was performed using whole cell lysate from liquid cultures of respective *S. oneidensis* strains labeled at the top of each lane. Also labeled are relevant bands in protein ladder (80 kDa, 58 kDa), as well as the approximate positions of proteins of interest (MtrC, OmcA) and fumarate reductase (FccA) which is present in all samples. Lanes 4 and 5 in both gels contain a dark band confirming the presence of heme associated with protein of interest (AP-tagged MtrC or OmcA). Wild type sample (Lane 1) was used as a positive control; respective gene deletion mutants Δ*mtrC* and Δ*omcA* (Lanes 2 and 3) are included as negative controls missing protein of interest MtrC or OmcA; and included as a secondary negative control is a cytochrome mutant (ΔMtr/Δ*mtrB*/Δ*mtrE*, Lane 6) missing a total of 8 periplasmic and outer membrane-associated cytochromes (including proteins of interest MtrC and OmcA).

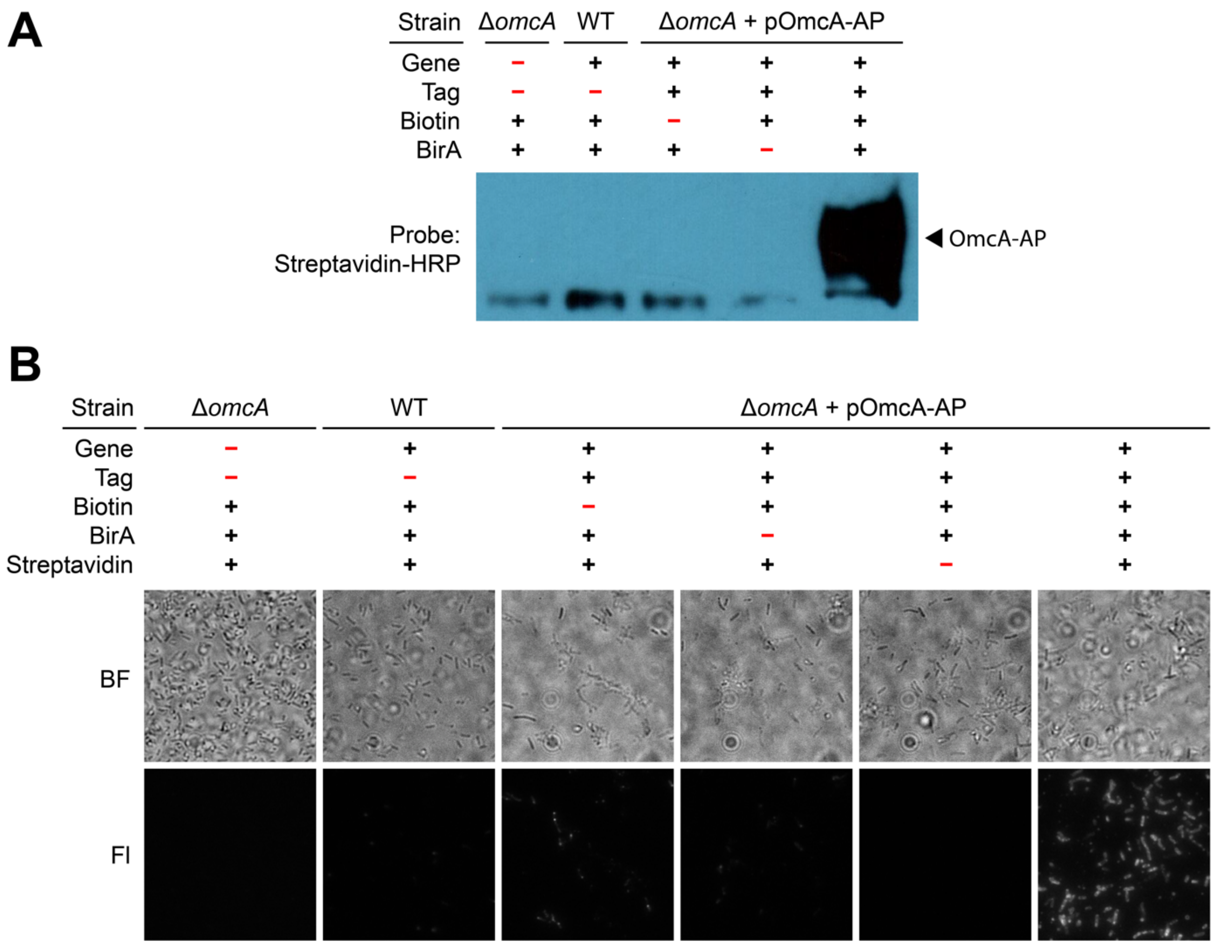


**Fig. S4. Key labeling controls demonstrate successful and specific labeling of OmcA. (*A*)** Western blot labeling control for OmcA where key parts of the labeling process were systematically omitted. When using streptavidin (streptavidin-horseradish peroxidase, HRP) to probe for biotinylated proteins, a thick dark band of biotinylated OmcA-AP is detected only in Lane 5 when all key components are present. The faint band slightly below labeled OmcA-AP (approx. 87 kDa) and present in all samples is an endogenously biotinylated protein (acetyl-CoA carboxylase, approx. 76 kDa) (22). **(*B*)** Microscopy labeling control for OmcA where key parts of the labeling process were systematically omitted. Top row contains brightfield (BF) images showing many cells in each sample. Bottom row images show fluorescence (Fl) signal from streptavidin-conjugated Alexa Fluor 647 that was used to detect biotinylated OmcA-AP; fluorescence labeling was detected strongly in the bottom right image, and only when all key labeling components were present. All microscopy images are 36.5 µm by 36.5 µm.

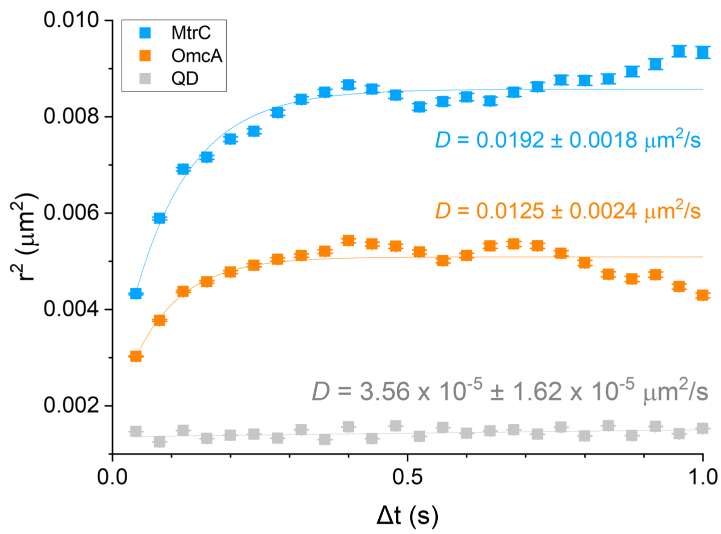


**Fig. S5. Apparent diffusion of quantum dot (QD) probes is distinct from QD-labeled MtrC and OmcA.** Ensemble mean squared displacement (MSD) analysis of QDs (gray), which were imaged on a coverslip in cell-free conditions. Here, Y-axis shows mean displacement squared ($r^2$) for each time lag ($\Delta t$) on the X-axis. Fitting this curve with a free (Brownian) diffusion model (Eqn. 3, gray fit line) yields the diffusion coefficient *D* for QDs, as labeled. This curve pools data from 56,835 trajectories from >1000 QDs. Error bars show $\pm\frac{r^2}{\sqrt{N}}$, where *N* is the number of independent data points (i.e., displacements) analyzed for a given $\Delta t$. For reference, ensemble MSD curves and confined diffusion fit for MtrC (blue) and OmcA (orange) on the cell surface are also included here, reproduced from Fig. 4.

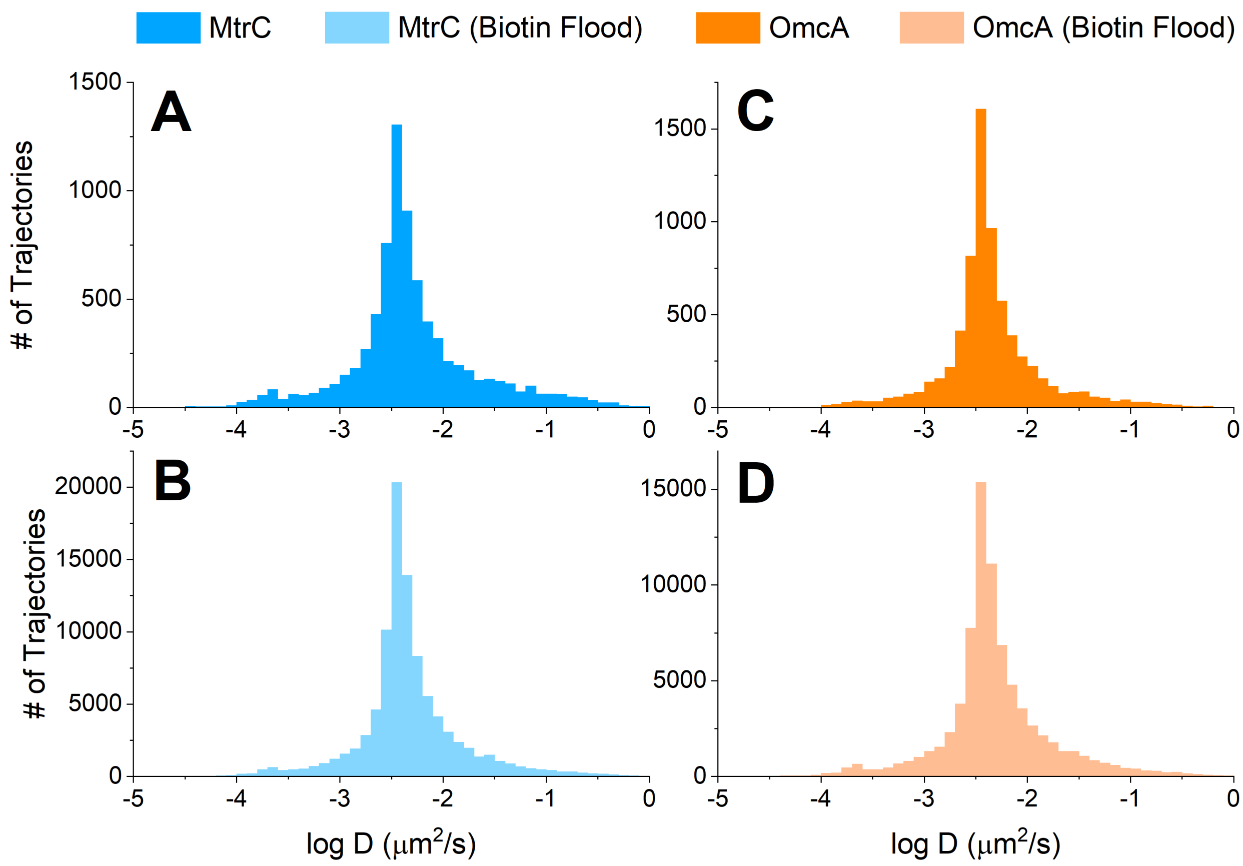


**Fig. S6. Diffusion is not limited by multivalent streptavidin quantum dot (QD) probes binding to multiple protein targets. (*A,C*)** Distribution of diffusion coefficients for QD-labeled MtrC-AP or OmcA-AP on the cell surface. These histograms represent 7,678 MtrC-AP and 7,109 OmcA-AP trajectories from 500-1,000 cells each. **(*B,D*)** Distribution of diffusion coefficients for QD-labeled MtrC-AP or OmcA-AP on the cell surface, where samples were saturated with excess biotin immediately after the QD labeling step. These histograms represent 96,163 MtrC-AP and 79,286 OmcA-AP trajectories, from >1,000 cells each. Each plot shows the distribution of diffusion coefficients for all individual trajectories in that dataset, calculated by fitting the free diffusion model (Eqn. 3) to the first 3 $\Delta t$ of each trajectory's mean squared displacement (MSD) curve.

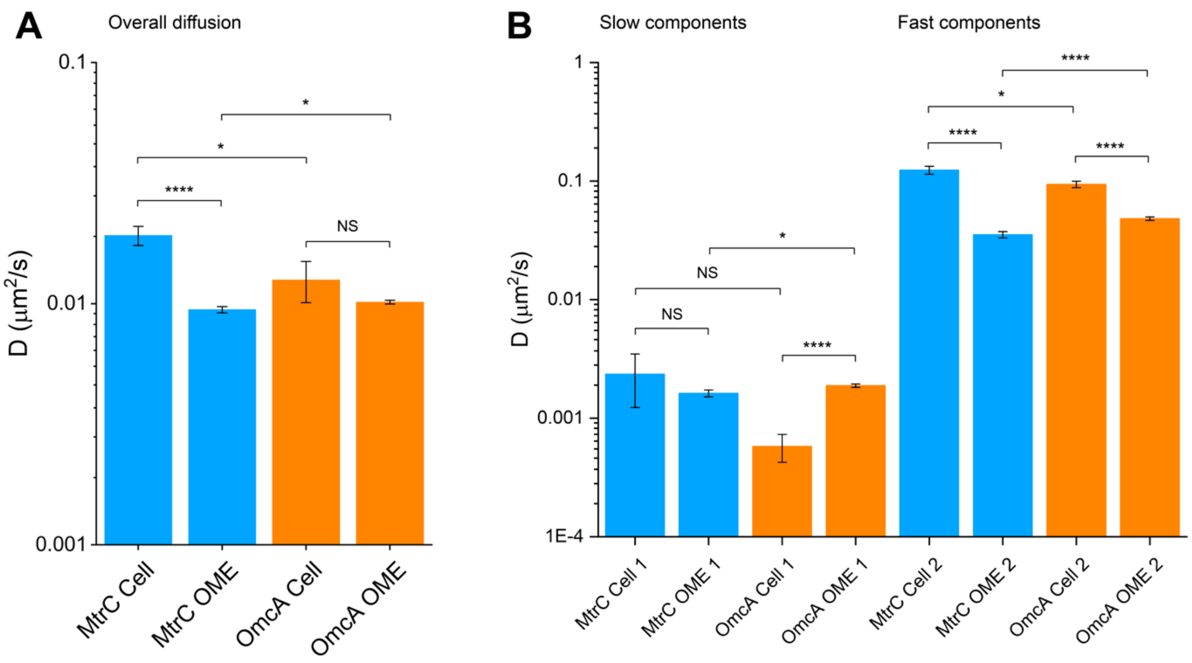


**Figure S7. Comparison of diffusion coefficients obtained in this study. (*A*)** Overall diffusion coefficients (± SD) for MtrC and OmcA on the cell surface or outer membrane extensions (OME). **(*B*)** Diffusion coefficients (± SD) for both fast and slow components of diffusion for MtrC and OmcA on the cell surface or on OMEs. (F-test, *$p < 0.05$, ****$p < 0.0001$, NS: not significant).

**Table S1.** Primers used in this study.

| Name | Sequence (5' to 3') | Description |
|---|---|---|
| MtrC Forward | GGAACCTCGAGGGGAATTCTATTTCCAGCATCC | 5-bp protection bases, XhoI cut site, binds to region upstream of MtrC |
| MtrC Reverse 1 | GATATCGTTCAGGCCTGAACCCATTTTCACTTTAGTGTGATC | Binds region of MtrC gene just before stop codon, adds first part of linker and AP tag sequence |
| OmcA Forward | GGAACCTCGAGGACTTAGTTAGCCTTACAGGTG | 5-bp protection bases, XhoI cut site, binds to region upstream of OmcA |
| OmcA Reverse 1 | GATATCGTTCAGGCCTGAACCGTTACCGTGTGCTTCCATCA | Binds region of OmcA gene just before stop codon, adds linker and first part of the AP tag sequence |
| MtrC/OmcA Reverse 2 | ACTCGATCTTCTGGGCCTCGAAGATATCGTTCAGGCCTGAAC | Binds to last 20-bp added by Reverse 1 primers in Round 1 of PCR extension; adds more of the AP tag sequence |
| MtrC/OmcA Reverse 3 | GGAACTCTAGATTACTCGTGCCACTCGATCTTCTGGGCCTCG | Binds to last 20-bp added by Reverse 2 primers in Round 2 of PCR extension; adds the rest of the AP tag sequence, stop codon, XbaI cut site, and 5-bp protection bases |
| M13 Forward (-20) | GTA AAA CGA CGG CCA GT | Used to check success of cloning and transformation by PCR gel and DNA sequencing |
| M13 Reverse (-27) | CAG GAA ACA GCT ATG AC | Used to check success of cloning and transformation by PCR gel and DNA sequencing |

**Table S2. Ingredients used for overhang PCR.** Volume is for a 25 µL reaction. For specific primers used in each step, see Table S4.

| PCR Ingredients | **Vol (µL)** | **Final Conc** |
|---|---|---|
| RNAse Free Water | *15.75* | |
| 5X Phusion HF Buffer | 5 | 1X |
| 10 mM dNTPs | 0.5 | 200 µM |
| F primer, 10 µM stock | 1.25 | 0.5 µM |
| R primer, 10 µM stock | 1.25 | 0.5 µM |
| Template DNA | 1 | ~45 ng |
| Phusion DNA Polymerase | 0.25 | 1 unit/50 µL rxn |

**Table S3. Thermocycler protocol for overhang PCR.** For specific annealing temperatures used for each primer pair, see Table S4.

| | Temp (°C) | Time | Cycles |
|---|---|---|---|
| Initial Denaturation | 98 | 30 s | |
| Denaturation | 98 | 10 s | x 5 |
| Annealing 1 | * | 30 s | |
| Extension | 72 | 1 min 15 s | |
| Denaturation | 98 | 10 s | x 30 |
| Annealing 2 | * | 30 s | |
| Extension | 72 | 1 min 15 s | |
| Final Extension | 72 | 7 min | |
| Hold | 4 | infinity | |

**Table S4. Specific primers and annealing temperatures used in overhang PCR.** Annealing temperatures were chosen for template-binding portions of each primer (Annealing 1) or for the full primer set (Annealing 2) according to the polymerase manufacturer's website (http://tmcalculator.neb.com/#!/main).

| | **F Primer** | **R Primer** | **Annealing 1 (°C)** | **Annealing 2 (°C)** |
|---|---|---|---|---|
| MtrC Round 1 | MtrC Forward | MtrC Reverse 1 | 57 | 72 |
| MtrC Round 2 | MtrC Forward | MtrC/OmcA Reverse 2 | 61 | 72 |
| MtrC Round 3 | MtrC Forward | MtrC/OmcA Reverse 3 | 64 | 72 |
| OmcA Round 1 | OmcA Forward | OmcA Reverse 1 | 56 | 72 |
| OmcA Round 2 | OmcA Forward | MtrC/OmcA Reverse 2 | 56 | 72 |
| OmcA Round 3 | OmcA Forward | MtrC/OmcA Reverse 3 | 56 | 72 |

**Table S5. Ingredients used for colony PCR.** Volume is for a 50 µL reaction. For colony PCR, MtrC or OmcA Forward primers were used along with MtrC/OmcA Reverse 1 primer to amplify DNA from putatively transformed *E. coli* DH5α colonies.

| | **Vol (µL)** | **Final Conc** |
|---|---|---|
| RNAse Free Water | *34.75* | |
| 5X OneTaq Reaction Buffer | 10 | 1X |
| 10 mM dNTPs | 1 | 200 µM |
| F primer, 10 µM stock | 1 | 0.5 µM |
| R primer, 10 µM stock | 1 | 0.5 µM |
| Template DNA | 2 | variable |
| OneTaq Quick-Load DNA Polymerase | 0.25 | 1.25 unit/50 µL rxn |

**Table S6. Thermocycler protocol for colony PCR.** Annealing temperatures were chosen for template-binding portions of each primer (Annealing 1) or for the full primer set (Annealing 2) according to the polymerase manufacturer's website (http://tmcalculator.neb.com/#!/main).

| | Temp ($^oC$) | Time | Cycles |
|---|---|---|---|
| Initial Denaturation | 94 | 30 s | |
| Denaturation | 94 | 30 s | x 5 |
| Annealing 1 | 44 (MtrC) or 50 (OmcA) | 60 s | |
| Extension | 68 | 2 min 30 s | |
| Denaturation | 94 | 30 s | x 25 |
| Annealing 2 | 62 | 60 s | |
| Extension | 68 | 2 min 30 s | |
| Final Extension | 68 | 5 min | |
| Hold | 4 | infinity | |

**Movie S1. Video and trajectories of a single quantum dot labeled OmcA-AP as it moves along the surface of a single cell.** Streptavidin-coated QD705 was used to detect exogenously biotinylated OmcA-AP. Here, the quantum dot signal (red) and its trajectories (white) are overlaid onto the mean intensity projection image of the cell labeled by lipid membrane dye FM 1-43FX (cyan). Video is from 85.64 s of time-lapse TIRF microscopy (40 ms/frame). This movie corresponds to Fig. 3D, which shows a snapshot of all trajectories in this video. Scale bar: 500 nm.

**Movie S2. Video of several quantum dot labeled OmcA-AP and their trajectories as they move along an outer membrane extension that appears to connect two cells.** Streptavidin-coated QD705 was used to detect exogenously biotinylated OmcA-AP. Here, the quantum dot signal (red) and its trajectories (white) are overlaid onto the mean intensity projection image of cell membrane and membrane extensions, labeled by lipid membrane dye FM 1-43FX (cyan). The movement of the quantum dots along the membrane extension is tracked over a short period (12 s) of time-lapse TIRF microscopy (40 ms/frame). This movie corresponds to Fig. 3E, which shows a snapshot of all trajectories from the full-length original video. Scale bar: 1 µm.

**Movie S3. Video of a quantum dot labeled MtrC-AP and its trajectories as it moves along an outer membrane extension.** Streptavidin-coated QD705 was used to detect exogenously biotinylated MtrC-AP. Here, the quantum dot signal (red) and its trajectories (white) are overlaid onto the mean intensity projection image of cell membrane and membrane extensions, labeled by lipid membrane dye FM 1-43FX (cyan). The movement of the quantum dots along the membrane extension is tracked over a short period (12.44 s) of time-lapse TIRF microscopy (40 ms/frame). Scale bar: 500 nm.

**Movie S4. Short simulation of electron transport (ET) assuming fast physical diffusion of redox carriers across the surface of a 190 nm long, 60 nm diameter membrane extension.** The surface is approximated as a 2D lattice array containing reduced redox carriers (red rectangles) and oxidized redox carriers (yellow rectangles). Electron transport starting from the top row of the lattice is simulated as redox carriers diffuse randomly in any direction to unoccupied positions in the lattice, and electron hopping occurs randomly in any direction between reduced to adjacent oxidized redox carriers. In this example, the fractional loading of redox carriers on the surface $X$ = 0.1. The ratio of hopping to diffusion timescale ($t_e/t_p$) is 300, to illustrate fast diffusion. Total length of time simulated in this movie is $3.3 \times 10^{-5}$ s.

**Movie S5. Short simulation of electron transport (ET) assuming slow physical diffusion of redox carriers across the surface of a 190 nm long, 60 nm diameter membrane extension.** The surface is approximated as a 2D lattice array containing reduced redox carriers (red rectangles) and oxidized redox carriers (yellow rectangles). Electron transport starting from the top row of the lattice is simulated as redox carriers diffuse randomly in any direction to unoccupied positions in the lattice, and electron hopping occurs randomly in any direction between reduced to adjacent oxidized redox carriers. In this example, the fractional loading of redox carriers on the surface $X$ = 0.1. The ratio of hopping to diffusion timescale ($t_e/t_p$) is 0.033, to illustrate slow diffusion. Total length of time simulated in this movie is 0.01 s.
